# Structural basis of unidirectional DNA recombination by the ϕC31 serine integrase

**DOI:** 10.1101/2025.05.02.651858

**Authors:** Y. Eugene Sun, Louie Aspinall, Agnel P. Joseph, Sean D. Colloms, W. Marshall Stark, Laura Spagnolo

## Abstract

Large serine integrases catalyse the integration and excision of bacteriophage (phage) DNA genomes into and from the genomes of their bacterial hosts by site-specific recombination^1^. These recombination reactions are notable for their unidirectionality: integrase recognises and efficiently recombines short (40–50-bp) sequences in the phage (*attP*) and bacterial (*attB*) genomes, but is inactive on the product sites *attL* and *attR* flanking the inserted prophage, ensuring stable integration. The reverse reaction (excision) only occurs in the presence of a second phage-encoded protein, the Recombination Directionality Factor (RDF). Their strict recombination directionality has made serine integrases versatile tools in emerging genome-editing technologies^2-5^; however, the structural basis of directionality remains. Here we report structures of ϕC31 integrase, the most-studied and most widely used member of the serine integrase family, in complexes with its DNA recombination sites, with and without its RDF. These structures correspond to four key mechanism steps: integration of the phage DNA genome (*attP* × *attB*); excision of the integrated prophage DNA, mediated by integrase and RDF (*attL* × *attR*); inhibition of integrase-catalysed excision in the absence of RDF (*attL* dimer complex); and inhibition of integrase-catalysed integration in the presence of RDF (*attB* dimer complex). Our data provide a mechanistic understanding of how the serine integrase recombination system establishes and regulates phage lysogeny, and lay the foundation for future development of integrase-based genome-editing technologies.

---

Many phages can establish a lysogenic phase by integrating their circular double-stranded DNA genomes into the genomes of their bacterial hosts. Integration, and the reverse process, excision, which re-establishes the lytic phase, are both catalysed by a phage-encoded site-specific recombinase enzyme called integrase. Integrase acts on ’attachment sites’ in the phage and bacterial DNA genomes (*attP* and *attB* respectively), and on the product sites *attL* and *attR* flanking the integrated prophage DNA. The *Streptomyces* phage ϕC31 integrase is one of a large group of these enzymes known as the serine integrases, which are characterised by a catalytic domain belonging to the serine recombinase superfamily^1,6^.

The ϕC31 site-specific recombination system has been studied extensively, *in vivo* and *in vitro^1^*. In common with other well-studied serine integrase systems, ϕC31 *attP* and *attB* sites (∼50 bp and ∼40 bp respectively) each bind an integrase dimer. The dimer-bound *att* sites interact via protein-protein interactions to form a synaptic complex in which *attP* and *attB* are held together on the outside of an integrase tetramer. Serine integrases are thought to share a ’subunit rotation’ strand exchange mechanism with all other serine recombinases^1,6^. It is proposed that integrase promotes double-strand cleavage of both synapsed *att* sites at specific phosphodiesters near their centres, using an active site serine residue. One half of the synaptic complex then rotates through 180° relative to the other half, and the cleavage steps are reversed to form recombinant sites (Fig. 1a). The *attB* × *attP* reaction, which requires only integrase, is very efficient and unidirectional. The product *attL* and *attR* sites do not recombine with each other, nor do any other pairs of *att* sites. Remarkably however, the presence of the phage-encoded RDF protein transforms the activity of integrase so that it promotes efficient *attL* × *attR* recombination, and *attB* × *attP* recombination is inhibited^7,8^. The structures of the serine integrase-associated RDFs including that of ϕC31 (also known as gp3), and the mechanism by which they switch recombination directionality, have remained unknown. These proteins show very high variability in size and amino acid sequence, with no obvious conserved features that could be correlated with their mechanism of action (refs. ^1,9^; see also below).

**Fig. 1:**
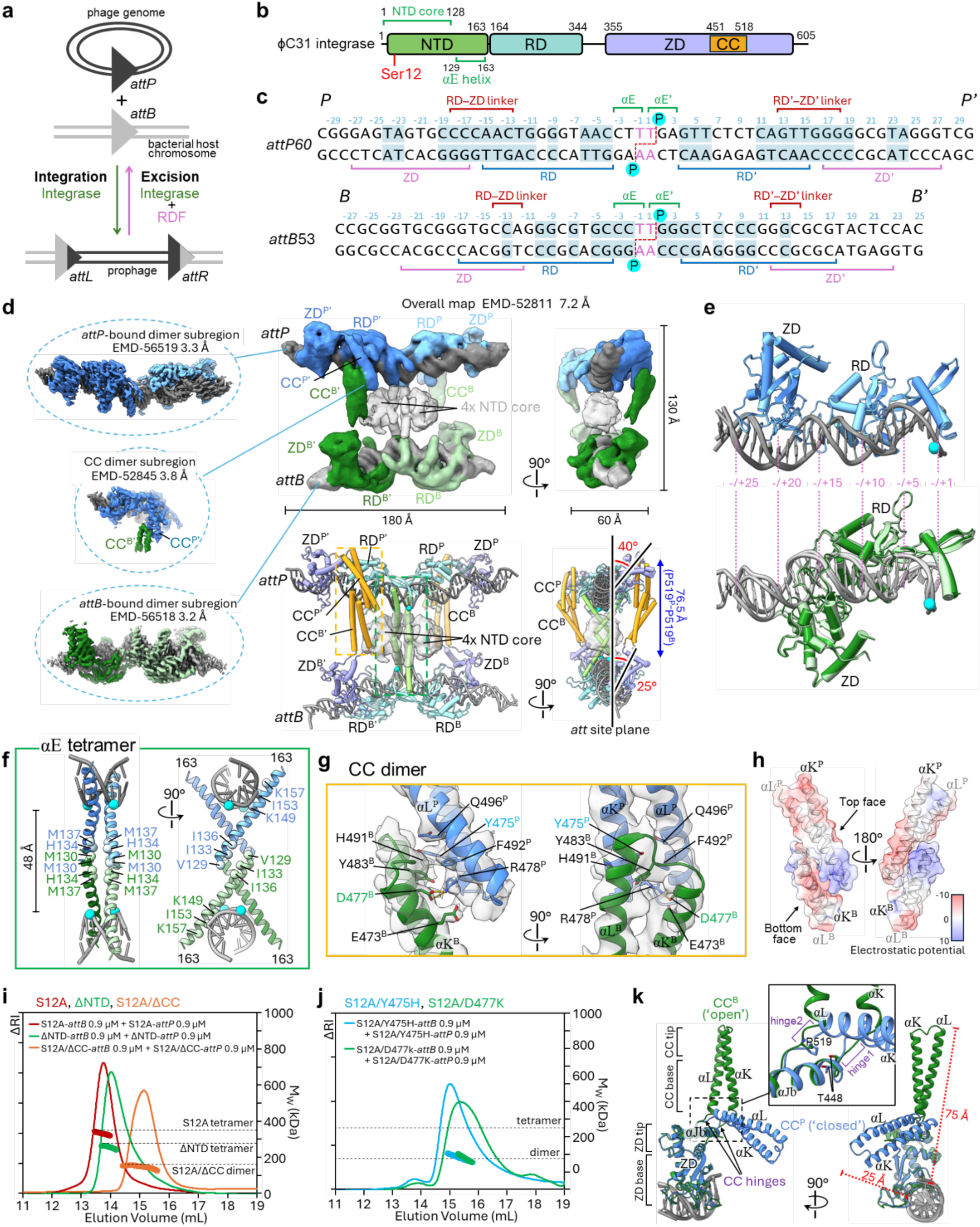
Structure of the ϕC31 integrase–*attP*–*attB* synaptic complex. **a**, Schematic of serine integrase-mediated integration excision during phage lysogeny. **b**, Domain architecture of the ϕC31 integrase. NTD: N-terminal catalytic domain; NTD core: conserved lobe in NTD that contains the catalytic serine (Ser12, marked in red); αE, conserved helix implicated in DNA binding and subunit rotation; RD, recombinase domain; ZD, zinc-ribbon domain; CC, coiled-coil motif. **c**, The 53-bp *attB*53 and 60-bp *attP*60 substrates used in cryo-EM experiments. The central 2 bp of each *att* site are highlighted in pink. The half-sites are denoted as *P*, *P’* (*attP*) and *B*, *B’* (*attB*). Sequences conserved in pairs of half-sites are highlighted with blue shading. Base-pair positions are numbered from the centre, with negative values for *B* and *P*, and positive values for *B’* and *P’*. Positions of the scissile phosphates are marked with cyan circles. Dashed red lines indicate the resulting 2 bp staggered ends after DNA cleavage. Solid brackets indicate the base-pair positions contacted by each integrase domain or motif, as observed in the cryo-EM structures. **d**, Cryo-electron microscopy (Cryo-EM) structure of the ϕC31 integrase^S12A^–*attP*–*attB* synaptic complex, showing two orthogonal views of the overall map (7.2 Å) and atomic model, and three high-resolution (3.2–3.8 Å) focused maps of local subregions (dashed circles) (see Extended Data Fig. 1 and Methods for details). *attB* and *attP* are coloured light and dark grey, respectively, in both the maps and models. In the cryo-EM maps, integrase is coloured by subunit, whereas in the models it is coloured by domain features (as in **b**). Phosphorus atoms of the four scissile phosphates are shown as cyan spheres. Density corresponding to the NTD tetramer, including the unmodelled regions (residues 1–128), is rendered as a grey transparent surface. The distance bridged by the CC dimer, measured between the CC anchor points (Cα of P519), is indicated in blue. The angles at which ZDs from a rotating dimer emerge from the *att* site plane are indicated in red, measured using a ZD axis passing through its base region near the bound DNA at R386 and tip region where CC protrudes at P519. Close-up views of the regions in yellow-and green-dashed boxes are shown in **f** and **g**. **e**, Comparisons of DNA-binding at each half-site; The *P* and *P’* (top), and *B* and *B’* (bottom) half-site structures are superimposed. Models are coloured by subunit as in the overall map in **d**. Similar orientations of *attP* and *attB* DNA are shown, with the site centres on the right and base-pair positions indicated. **f**, Close-up views of the NTD αE helix tetramer and bound DNA. Selected residues involved in the tetramer interface and DNA binding have their Cα atoms shown as spheres. **g**, Close-up views of the CC motif dimer interface, showing the model from focused refinement (PDB: 9IFC) with the corresponding cryo-EM density (EMD-52845) overlaid as a transparent surface. Key side-chains engaged in contacts between the motifs are shown as sticks. Residues labelled in blue and green were targeted by mutagenesis in **j**. **h**, Views of the CC motif dimer interface; transparent cryo-EM density surface is coloured by electrostatic potential. Black arrows mark the ‘top’ and ‘bottom’ faces of CC^P^ and CC^B^, respectively. The left panel in **h** shows a similar view to the left panel in **g**. **i**,**j**, SEC-MALLS analysis of the integrase–*att* complexes formed by full-length S12A integrase and its truncation mutants (**i**), and point mutants (**j**). The protein elution profile is shown as elution volume (x-axis) against differential refractive index (ΔRI) (y-axis). Circular markers represent the calculated weight-averaged molar mass (M_W_). Black dashed lines mark the expected M_W_ values for the indicated dimer–*att(B or P)* and tetramer– *attB*–*attP* complexes. ΔNTD, residues 1–140 deleted; S12A/ΔCC, residues 450–515 deleted. **k**, Structural comparison of CC motif trajectories in monomers bound to *attB* and *attP*. Structures are superimposed at their ZD Cα atoms (excluding residues in the CC motifs). Black arrows mark the flexible regions in the αJb–αK and αL–αM loops that act as hinges between the CC motifs and the remainder of the ZDs. The inset shows a close-up of these hinges, highlighting residues T448 and P519 at the edge of the ZD, whose positions remain relatively invariant and thus serve as ‘CC anchor points’. Distances from the CC motif tip (Cα of G485) and the *att* site DNA are shown in red. All structural displays were generated using ChimeraX^35^ version 1.10.1.

The catalytic domain of ϕC31 integrase (NTD, residues 1–163) is at the N-terminal end of the 605-amino acid polypeptide chain. The NTD is followed by a central ’recombinase domain’ (RD, residues 164–344, also known as DNA-binding domain 1), and finally a large C-terminal ’zinc ribbon domain’ (ZD, residues 355–605, also known as DNA-binding domain 2) (Fig. 1b). The RD and ZD are involved in specific recognition of the *attB* and *attP* recombination sites. Within the ZD is a coiled-coil structural motif (CC, residues 451–518) which plays crucial roles in synapsis and directionality control^1,10-12^. Previous structural and biochemical studies have led to a model in which directionality is established through *att* site-specific integrase conformations that selectively permit certain *att* sites to assemble into synaptic complexes^13-16^. However, to date there are no high-resolution structures of any full-length serine integrase in complexes with its sites, nor of synaptic intermediates or RDF–integrase complexes.

Here, we present high-resolution cryo-EM structures of full-length ϕC31 integrase protein in complexes with its recombination sites, with and without RDF. These structures reveal how reactivity is governed by the trajectories of the CC motifs as they emerge from the individual integrase subunits bound at the *att* sites. We observe a synaptic complex comprising *attB* and *attP* sites bound together by an integrase tetramer, the intermediate immediately prior to ’integrative’ DNA cleavage and recombination. However, on the product *attL* site, integrase binds to form a locked/inactive dimer complex in the absence of RDF; but in the presence of RDF, we observe instead a synaptic complex comprising two *attL* sites and an integrase tetramer, with each integrase subunit bound to an RDF subunit. This corresponds to the intermediate expected immediately prior to ’excisive’ (*attL* × *attR*) recombination, a second *attL* site acting as a surrogate for *attR*. Finally, we observe a locked/inactive dimer complex of an RDF-bound integrase dimer with *attB*, revealing how the RDF inhibits *attB* × *attP* recombination. The synaptic complexes are stabilised by interactions of the CC motifs bound at one site with those bound at the other (inter-site), whereas in the locked dimer complexes the two CC motifs from subunits bound to the same *att* site interact (intra-site). Our data allow us to account for directionality of integrase-mediated recombination in the absence and presence of RDF.

## Results

### Structure of the attB–attP synaptic complex

We reconstituted the ϕC31 integrase–*attB*–*attP* synaptic complex, using a 53-bp *attB* and a 60-bp *attP* duplex together with a cleavage-deficient S12A integrase mutant^13^ (Fig. 1b,c).The complex is thus trapped in a pre-cleavage state. Purified complexes were analysed by single-particle cryo-EM, producing a 7.2 Å reconstruction of the full integrase–*attB*–*attP* complex (Fig. 1d (right top)). Global high-resolution reconstruction was precluded by pronounced regional motion (Extended Data Fig. 1a). Focused classification and refinement^17^ of the mobile subregions were applied to improve map quality, yielding well-resolved reconstructions of the *attB*-and *attP*-bound dimer regions, and a third region comprising an *attP*-bound monomer with a well-defined interface between its CC motif and that of its partner *attB*-bound integrase subunit, at resolutions of 3.2 Å, 3.3 Å, and 3.8 Å, respectively (Fig. 1d (left) and Extended Data Fig. 1). Atomic models were built for both *att* site duplexes, and all four integrase monomers, apart from residues 1–128 and 590–605 (see Methods).

The structure of the *attB*–*attP* synaptic complex reveals distinct integrase conformations at *attB* and *attP*, particularly in the positioning of the ZDs on the DNA and the trajectories of the CC motifs that extend from them (Fig. 1d,e and Extended Data Fig. 2a). The two *att* site duplexes are ∼50 Å apart at the centre of the complex, with the two pairs of scissile phosphodiesters facing each other. Each duplex bends away from the centre, so that the ends of *attB* and *attP* are ∼80 Å apart. The long axes of the *att* sites lie nearly in the same plane and are almost parallel. The complex exhibits imperfect twofold rotational symmetry, with the symmetry axis passing through the central two base-pairs of the two *att* sites. The RD and ZD domains bind primarily in the major grooves, with the RDs close to the centres of the *att* sites and the ZDs near the ends.

There are several inter-subunit protein-protein interfaces between integrase monomers: a tetrameric assembly of αE helices near the central twofold axis of synaptic complex, and a pair of distally located CC–CC dimers. Although we did not build a model for the remainder of the catalytic domains, their diffuse density is seen surrounding the αE helices (shown in transparent grey in Fig. 1d).

The αE tetrameric assembly has 222 symmetry, consistent with that observed in tetramers of other serine recombinases^18-21^ (Fig. 1f and Extended Data Fig. 2b). Each αE (residues 129–163) forms a continuous 10-turn helix, with its C-terminal portions inserting into the minor groove at the central 6 bp of each *att* site. The face of αE that contacts DNA, comprising K149, I153, and K157, directs towards the scissile phosphate. The two helices of each ‘rotating dimer’ (that is, a pair of integrase subunits bound to the *B* and *P* half-sites, or the *B’* and *P’* half-sites, which are predicted to move together during subunit rotation) pack against one another using their two N-terminal helical turns (residues 129–136). This interface involves hydrophobic residues V129, I133, and I136 from each monomer, which are positioned on the opposite face of the helix from the DNA-binding face (Fig. 1f).

The CC motifs dimerise within each rotating dimer (as defined above), interacting in a ‘top-to-bottom’ arrangement, similar to that seen in the crystal structure of the isolated CC motif of LI integrase^11^ (Fig. 1d,g,h and Extended Data Fig. 2c–e). The dimer interface involves the tip region of the CC (residues 468–479), corresponding to the C-and N-terminal portions of αK and αL, respectively. These pack together into a four-helix bundle in an antiparallel arrangement, resulting in an extended dimer spanning a distance of 73.1 Å (measured from the base of each CC, Cα of E517) (Fig. 1g,h). In this arrangement, the tip region of each αK, which is rich in acidic residues, is packed against the highly basic αL region of the opposing monomer, ensuring top-to-bottom stacking (Fig. 1h). The interface also contains a salt-bridge between D477^B^ on the final turn of αL and R478^P^ on the opposing monomer (superscripts B and P indicate the CC motifs bound to *attB* and *attP* respectively), and a hydrophobic core involving Y475^P^, F492^P^, and Y483^B^ (Fig. 1g). We note that both the charged regions and the hydrophobic residues in the CC motif interface are highly conserved within the *Streptomyces* group of phage serine integrases (Extended Data Fig. 3).

To assess the individual contributions of these interfaces to synapsis, we generated two integrase deletion mutants; ΔNTD (residues 1–140 deleted), and S12A/ΔCC (residues 450–515 deleted). We then measured the ability of these mutants to form *attB*–*attP* synaptic complexes, using Size-Exclusion Chromatography coupled to Multi-Angle Laser Light Scattering (SEC-MALLS). A sample containing equal amounts of purified full-length S12A–*attB* complex and S12A–*attP* complexes (see Methods) eluted as a single peak with a M_W_ (weight-averaged molar mass) of 327 kDa, closely matching the expected mass of 348 kDa for the full tetramer–*attB*–*attP* synaptic complex. Similarly, the ΔNTD mutant readily formed synaptic complexes (Fig. 1i). In contrast, the measured M_W_ of the peak observed with the S12A/ΔCC mutant was consistent with the formation of dimer–*att* complexes. We conclude that the ΔCC mutant can bind *attB* and *attP* to form on-site dimer (or two-subunit) complexes, but these are unable to interact to form synaptic complexes. To confirm that this synapsis deficiency was due to inability to form the CC motif interface, we created two integrase point mutants, S12A/Y475H and S12A/D477K, substituting residues involved in the observed interface. These mutants also failed to promote synapsis (Fig. 1j). Taken together, these results show that the CC motif serves as the main driver of ϕC31 integrase-mediated synapsis.

Our structure reveals how the *att* site DNA sequences dictate selective *attB* × *attP* synapsis. At *attP*, integrase binding closely resembles that observed in the crystal structure of LI integrase C-terminal fragment bound to its *attP* half-site^15^. The RD and ZD bind to adjacent major grooves on the same face of the DNA (Fig. 1e and Extended Data Fig. 2a), at base-pair positions -/+4–15 and -/+17–27, respectively (Fig. 1c). The 10-residue ZD–RD linker (residues 345–354) binds the same DNA face as the RD and ZD, inserting into the minor groove at positions -/+13–18, a motif with identical sequence in the two *attP* half-sites (Figure 1c,e and Extended Data Fig. 2a). At *attB*, the RD binds each half-site at the same distance from the centre as at *attP*. However, the RD–ZD linker crosses the minor groove, contacting positions -/+12–14, and the adjacent ZD is shifted 5 bp closer to the site centre (positions -/+12–22), so that the ZDs bound at *attB* are on the opposite face of the DNA (relative to the centre and the RDs) from those bound at *attP* (Fig. 1e and Extended Data Fig. 2a). This differential ZD positioning generates a synapse in which the ZDs from each rotating dimer emerge from the plane of the DNA sites at similar angles (Fig. 1d (bottom right)). As a result, the tip region of ZD from which the CC protrudes (Fig. 1d,k), is unobstructed by DNA, enabling inter-site CC dimerization. Any substantially different ZD configurations, as would be present in *attP*–*attP* and *attB*–*attB* pairings, would dramatically alter the inter-ZD distance, likely disfavouring these pairings.

The CC motif has a flexible hinge that joins its αK and αL helices to the ZD, allowing CC^B^ and CC^P^ to adopt the very different trajectories that enable synapsis (Fig. 1k). Residue P519, in the ZD tip, is positioned just beyond the end of αL and remains essentially fixed relative to the main body of the ZD in all integrase subunit structures, thus providing a reliable reference point (the ‘CC anchor point’) for comparison of CC trajectories and RDF-induced rearrangements (see below). In the synaptic complex, the CC anchor points are spaced 76.5 Å apart (Fig. 1d). The CC^P^ motifs follow a trajectory that extends toward the centre of *attP* and remain near the *attP* DNA (Fig. 1k). In contrast, the CC^B^ motifs are nearly perpendicular to the *attB* DNA, and reach across the synapse to contact CC^P^. We term these distinct conformations of CC^P^ and CC^B^ ‘closed’ and ‘open’, respectively.

### Structure of the attL-bound dimer complex

Integration generates two new *att* sites, *attL* and *attR*, each of which has one half-site derived from *attB* and one from *attP* (Fig. 1a,c and 2a). Because both *attB* and *attP* have partial two-fold symmetry, *attL* and *attR* are structurally and biochemically similar, with the only functional distinction being the orientation of the central 2 bp that determines their identity^22,23^. In the absence of RDF, *attL* and *attR* cannot form synaptic complexes, either with themselves or with each other, thereby ensuring unidirectional integration^10,13,14^. To understand the basis of this synapsis inhibition, we determined the EM structure of the ϕC31 integrase dimer–*attL* complex (Fig. 2b, Extended Data Fig. 4 and Methods). Atomic models were built for the *attL* DNA together with nearly all the protein of each monomer, including the NTD core regions.

**Fig. 2:**
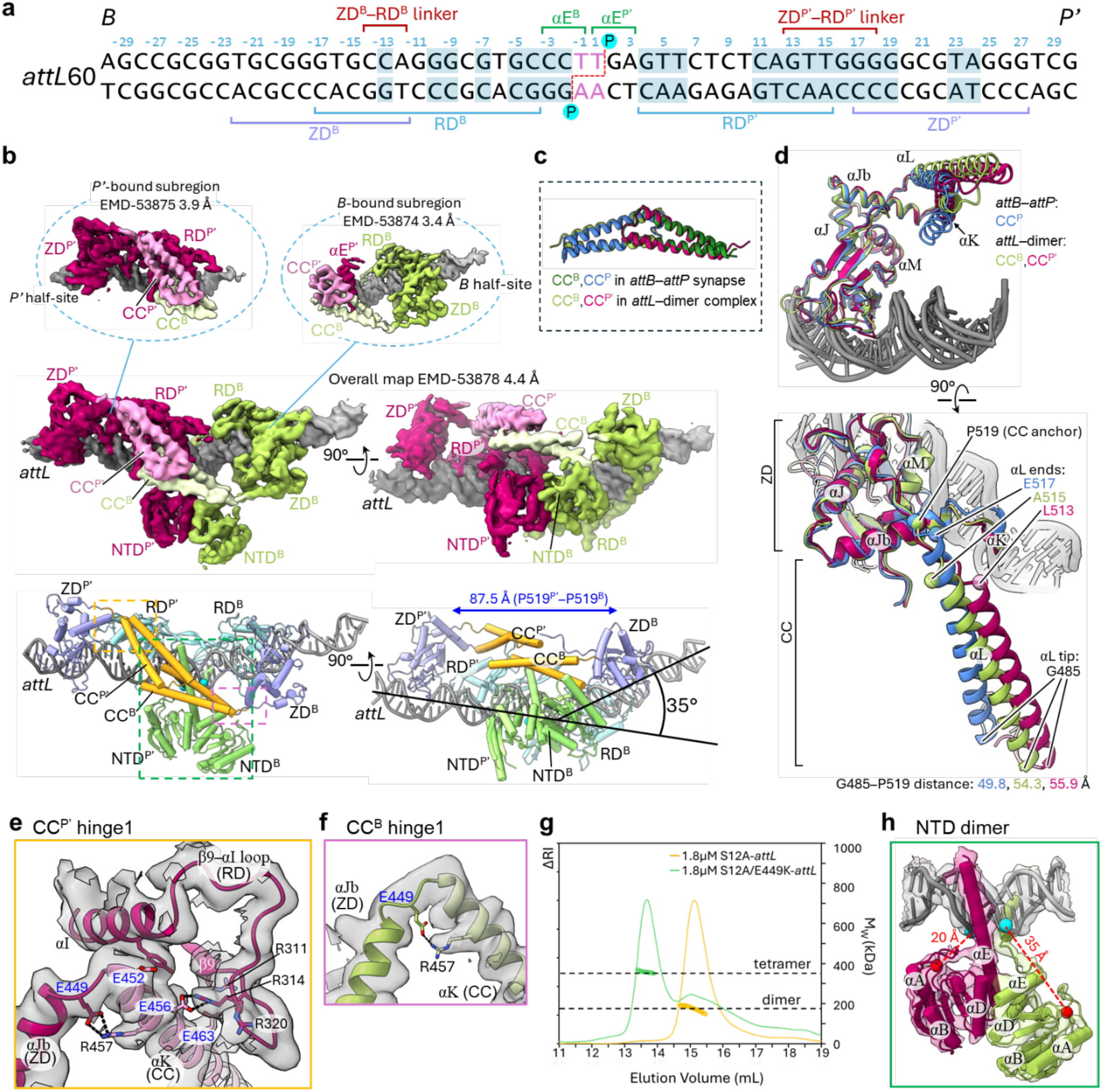
Structure of the integrase–*attL* dimer complex. **a**, Sequence of the *attL*60 DNA used in cryo-EM structural determination, with the same annotation and half-site highlighting as in Fig. 1c. **b**, Cryo-EM maps and the atomic model of the integrase–*attL* complex. High-resolution local maps are in dashed circles. *B* and *P’* half-sites of *attL* are coloured light and dark grey, respectively. Integrase is coloured by subunit (magenta and olive) in the maps and by domain features (as in Fig. 1b) in the models. Phosphorus atoms of the scissile phosphates are in cyan. The distance bridged by the CC dimer and the DNA bend angle are indicated in the bottom right panel. Close-up views of the regions in dashed boxes are shown in **e**,**f**,**h**. **c**,**d** Structural comparisons of the CC motif dimers (**c**) and trajectories (**d**) in the integrase–*attB*–*attP* synaptic complex and the integrase dimer–*attL* complex. For clarity, the CC^B^ in the synaptic complex (which adopts the open CC trajectory) is omitted in **d**. The αK helices in the bottom panels are also hidden for clarity. The reach of each CC, defined as the distance between the CC anchor point (P519) and the tip of αL (G485), is shown. **e**, Close-up view of the hinge1 region of CC^P’^, showing the connection of αJb and αK, and the nearby interface between αK and the β9–αI loop in RD^P’^. Acidic residues (blue labels) previously identified to confer hyperactive^10^ *attL* × *attR* recombination, and the basic residues they interact with, are shown as sticks. **f**, Close-up of hinge1 of CC^B^, showing the salt-bridge between E449 and R457. **g**, SEC-MALLS analysis of the *attL*-bound complexes made by the integrase^S12A^ and the S12A/E449K double mutant. Coloured lines and circular markers show the elution profiles and M_W_, respectively. **h**, Close-up view of the NTD dimer, with the cryo-EM density overlaid as transparent surface. The Cα atoms of the catalytic S12 residues are shown as red spheres, and the phosphorus atoms of the scissile phosphates as cyan spheres. Distances between the S12 residues and their respective scissile phosphates are indicated in red.

The *attL*–integrase dimer structure shows that monomers bound at the *B* and *P’* half-sites retain the same DNA-binding configurations as the corresponding monomers in the *attB*–*attP* synapse, so that the ZD domains are roughly on the same DNA face of *attL*. Both CC motifs extend towards the centre of the complex, where their tips make a ’top-to-bottom’ interface, nearly identical to that observed in the *attB*–*attP* synaptic complex. Interestingly however, the polarity of the interaction is switched, so that the top face of CC^P’^ is interacting with the bottom face of CC^B^, as defined above (Fig. 2c). The intra-site CC interaction is accompanied by altered hinge conformations; both CC motifs adopt a retracted trajectory resembling the ‘closed’ CC^P^ motifs in the *attB*–*attP* synaptic complex (Fig. 2d). The distance between the CC anchor points at the ZD tips is 87.5 Å, 11 Å longer than that observed for the *attB*–*attP* synaptic complex CC dimers (Compare Fig. 1d (bottom right) with Fig. 2b (bottom right)). This increase is accounted for by unwinding of the C-terminus of αL; In the *attL*-dimer complex, αL^B^ and αL^P’^ terminate at A515 and L513, respectively, whereas their counterparts in the *attB*–*attP* synapse end at E517 (Fig. 2d (bottom)). Bending of the *attL* DNA by ∼35° towards the interacting CC motifs also facilitates the CC interaction by bringing the CC anchor points closer together (Fig. 2b).

A network of electrostatic interactions near the hinge seems to stabilise the closed CC motif trajectories in the *attL*-bound structure. At both hinges, E449 at the αJb–αK junction forms a salt bridge with R457 in the second turn of αK (Fig. 2e,f). The first four turns of αK also create a highly acidic surface, consisting of E452, E456, and E463 which, for the *P*’-bound monomer, packs intramolecularly against the basic β9–αI loop in the RD, comprising R311, R314, and R320 (Fig. 2e). Strikingly, when each of these four glutamates was mutated to lysine, the mutant integrases were all shown to be ‘hyperactive’; that is, they promoted *attL* × *attR* recombination even in the absence of RDF^10^. Sequence analysis reveals that these charged glutamate and arginine residues are highly conserved within the *Streptomyces* group (Extended Data Fig. 3). To test the role of the E449–R457 interaction, we generated the integrase S12A/E449K mutant, and analysed its interactions with *attL* by SEC-MALLS. Unlike integrase S12A, the S12A/E449K double mutant readily produced *attL*–*attL* synaptic complexes (Fig. 2g). We therefore hypothesise that E449K and other activating mutations near the CC hinge relieve inhibition of integrase-mediated *attL* and/or *attR* synapsis, likely by destabilising the closed CC trajectories.

The NTDs in the *attL* complex pack together to form a dimer (Fig. 2h). The αE helix of NTD^P’^ is continuous, similar to that seen in the synaptic complex, whereas that of NTD^B^ is unwound at residues 140–148, allowing the NTDs to dimerise. The dimer interface involves αD and the N-terminal end of αE from each monomer, with a central hydrophobic core including the side chains of I103 on αD and I133 on αE (Fig. 2h; see also Fig. 4c). The NTDs appear to be locked in a cleavage-deficient conformation, with the active serines positioned 20 Å and 35 Å away from their respective scissile phosphates. Notably, both the NTD dimer interface and the conformation of the individual NTD monomers in the *attL*-bound structure are incompatible with the αE tetramer formed in the *attB*– *attP* synaptic complex, which is predicted to represent a ‘ready-to-cleave’ state (see Fig. 5a). We therefore propose that the transition from a cleavage-deficient dimer to a cleavage-competent tetramer involves the dissociation of the intra-site NTD dimer and a conformational change within each NTD monomer (see below).

Taken together, these results show that the ZD domains of integrase subunits bound to *attL* adopt conformations that permit intra-site CC motif dimerization, thereby locking *attL* in a synapsis-deficient state. Concurrently, the NTDs form a dimer in which each monomer is held in a cleavage-deficient conformation, preventing unwanted *att* site cleavage in the absence of synapsis. We infer that the structurally similar *attR* adopts comparable ZD binding positions, producing a similar locked dimer complex.

### Excisive synaptic complex structure

To understand how the ϕC31 RDF (gp3) promotes excision, we determined the structure of a synaptic complex consisting of two *attL* sites bound by four copies of a ϕC31 integrase-RDF fusion protein^24^. A 32-residue flexible linker was used to connect the C-terminus of the integrase S12A mutant to the N-terminus of the RDF protein (Extended Data Fig. 5a), in order to stabilise the complex for structural determination. This protein made a complex with *attL*, which had a molecular mass consistent with the predicted synaptic complex comprising two copies of *attL* and four fusion protein subunits (Extended Data Fig. 5b). The fusion protein complex was analysed by cryo-EM (Fig. 3a, Extended Data Fig. 6 and Methods). Atomic models were built for both *attL* duplexes, all four RDFs (residues 23–239), and integrase residues 129–592. The flexible fusion linker was not resolved. As in the *attB*–*attP* synaptic complex, densities for the NTD core regions were visible but insufficient for reliable modelling, and are therefore depicted as transparent grey in Fig. 3a.

**Fig. 3:**
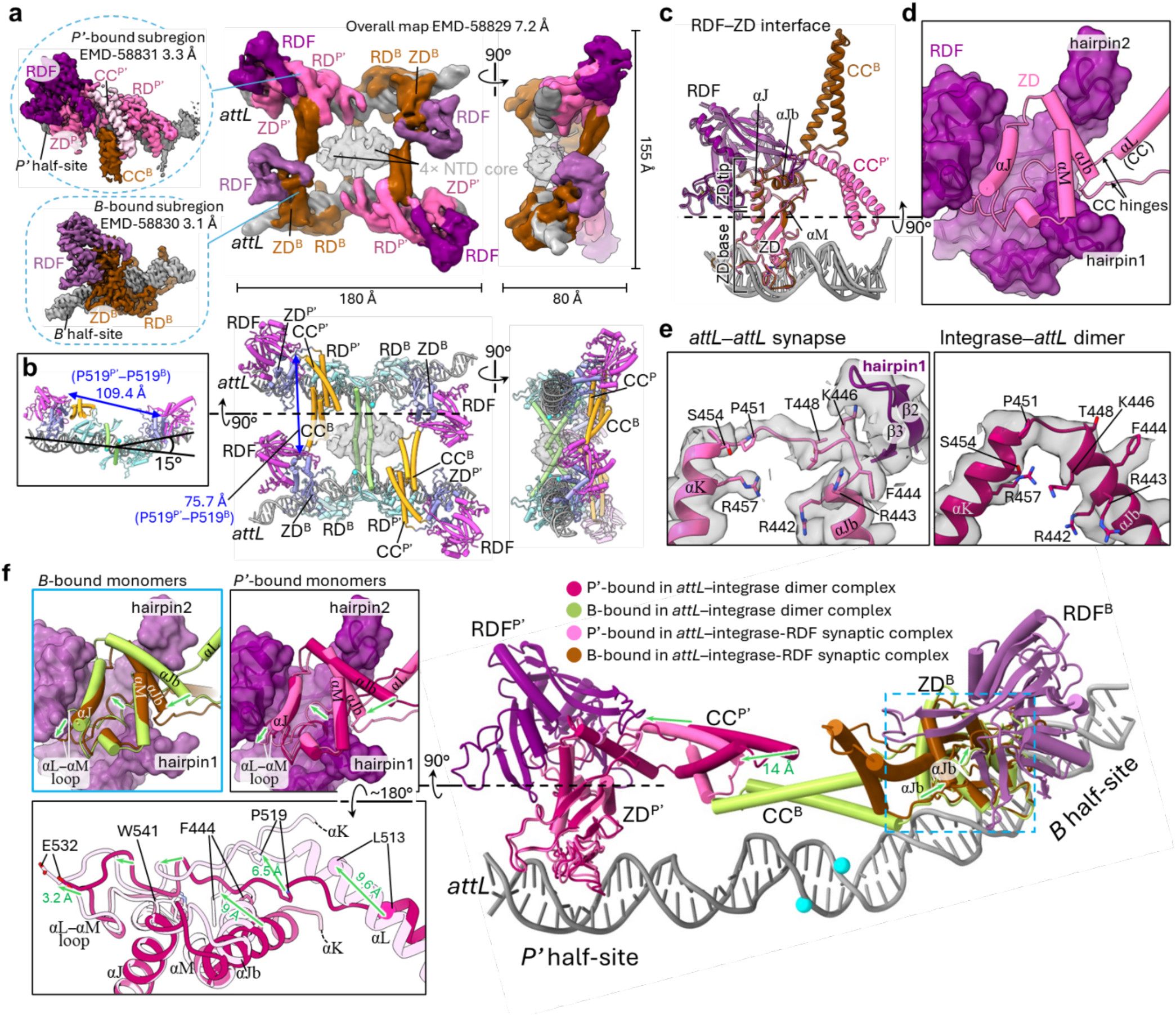
Structure of the integrase^S12A^-RDF–*attL* excisive synaptic complex. **a**,**b**, Cryo-EM maps and the atomic model of the integrase^S12A^-*attL*–RDF synaptic complex comprising two *attL* sites (grey) bound with four integrase^S12A^-RDF fusion proteins (coloured by subunits in the maps and by protein features in the models). High-resolution local maps are shown in dashed circles. Inset (**b**) shows a cross-sectional view. The distances between one pair of inter- and intra-site CC anchor points (P519) are indicated in blue. **c**, Structural comparisons of the ZD–RDF interaction at *B*- and *P’*-bound monomers. The structures are superimposed using Cα atoms in the RDF and ZD (excluding those in the CC motifs). **d**, Cross-sectional view showing the ZD^P^–RDF interface, with the ZD and the RDF shown as cartoon and surface view, respectively. **e**, Structural comparison of the hinge1 region of a *P’*-bound monomer in the integrase^S12A^–*attL*– RDF synaptic complex and the integrase dimer–*attL* complex. Cryo-EM maps are overlaid in transparent grey. **f**, Structural comparisons of monomers in the *attL*–integrase^S12A^-RDF and *attL*–integrase complexes, highlighting the RDF-induced confomational changes at the ZD tips and CC hinges. The panel on the right is an overview showing the *attL* DNA with two bound ZD domains (magenta and olive) from the *attL*–integrase structure. The ZD (pink and brown) and RDF (dark and light purple) regions of their RDF-bound counterparts in the *attL*–integrase^S12A^-RDF structure are overlaid using Cα atoms in the ZD (excluding those in the CC motifs). The panels on the left display close-up views at the ZD tip regions, showing the RDF-induced changes in the *B*- and *P’*-bound monomers, boxed in blue and black, respectively. For clarity, the RDF is hidden in the left bottom panel. Green arrows mark the trajectory shifts at key ZD and CC positions upon RDF binding, which involve displacements directed away from the *att* site centre where the CC motifs dimerise. Selected distances are indicated in green.

The structure of the RDF resembles a ‘three-pronged clamp’ that grasps onto the integrase ZD tip region, with very similar conformations at the *B* and *P’* half-sites of *attL* (Fig. 3c). A ∼20-Å-wide channel is formed between the larger prong (comprising αB and β8–10 of RDF) and two smaller prongs, each consisting of a β-hairpin. We refer to these as hairpin1 and hairpin2, which are formed by β2–3 and β12–13 respectively (Extended Data Fig. 5c). The channel accommodates the bulk of the ZD tip, including αJ, αJb, and the αL–αM loop. A gap between hairpin1 and hairpin2 allows passage of the CC motif hinges, from which the CC protrudes (Fig. 3d). The channel is enriched in basic residues that contact multiple acidic patches on the integrase ZD (Extended Data Fig. 5e,f). Additionally, hydrophobic residues on hairpin1 and hairpin2 contact the integrase CC hinges, engaging αJb and the αL–αM loop (Extended Data Fig. 5g,h). Collectively, these RDF–integrase contacts result in an extensive buried surface of ∼1600 Å^2^ (ref.^25^). Comparisons with RDF-free integrase subunit structures in the *attL*-dimer complex reveal substantial remodelling of the ZD tip upon RDF binding (Fig. 3f); in the RDF-bound structures, both CC anchor points (Cα of P519) are shifted away from the centre of *attL*, by 5.4 Å and 6.5 Å for the *B*-and *P’*-bound integrase subunit, respectively. In addition, the αL C-termini rewind to E517, further moving the CC tips away from the centre (Fig. 5f). We propose that the RDF binds to the integrase subunits of the *attL*-dimer complex to promote these changes, breaking the inhibitory CC dimer and thus freeing the CC motifs to make synaptic interactions.

Interestingly, another prominent RDF-induced remodelling of integrase occurs at the αJb–αK junction, where the helices are unwound, expanding the αJb–αK loop from three residues (E449– P451) to ten residues (G445–S454) (Fig. 3e). This rearrangement appears to be imposed by hairpin1, whose binding creates steric hindrance that prematurely terminates αJb.

### Structure of an attB–integrase–RDF complex

Inhibition of *attB* × *attP* recombination by RDF has been noted previously^7,8^, but the mechanism was unclear. We used SEC-MALLS to directly assess whether the RDF affects *attB*– *attP* synaptic complex formation (Fig. 4a). When RDF was added to pre-formed *attB*–*attP* synaptic complex, a new peak was observed with M_W_, closely matching that predicted for an integrase dimer– RDF–*att* complex. Indeed, SDS-PAGE analysis of this peak confirmed the presence of integrase and RDF, consistent with RDF-induced dissociation of the synaptic complex into dimeric assemblies (Extended Data Fig. 7).

**Fig. 4:**
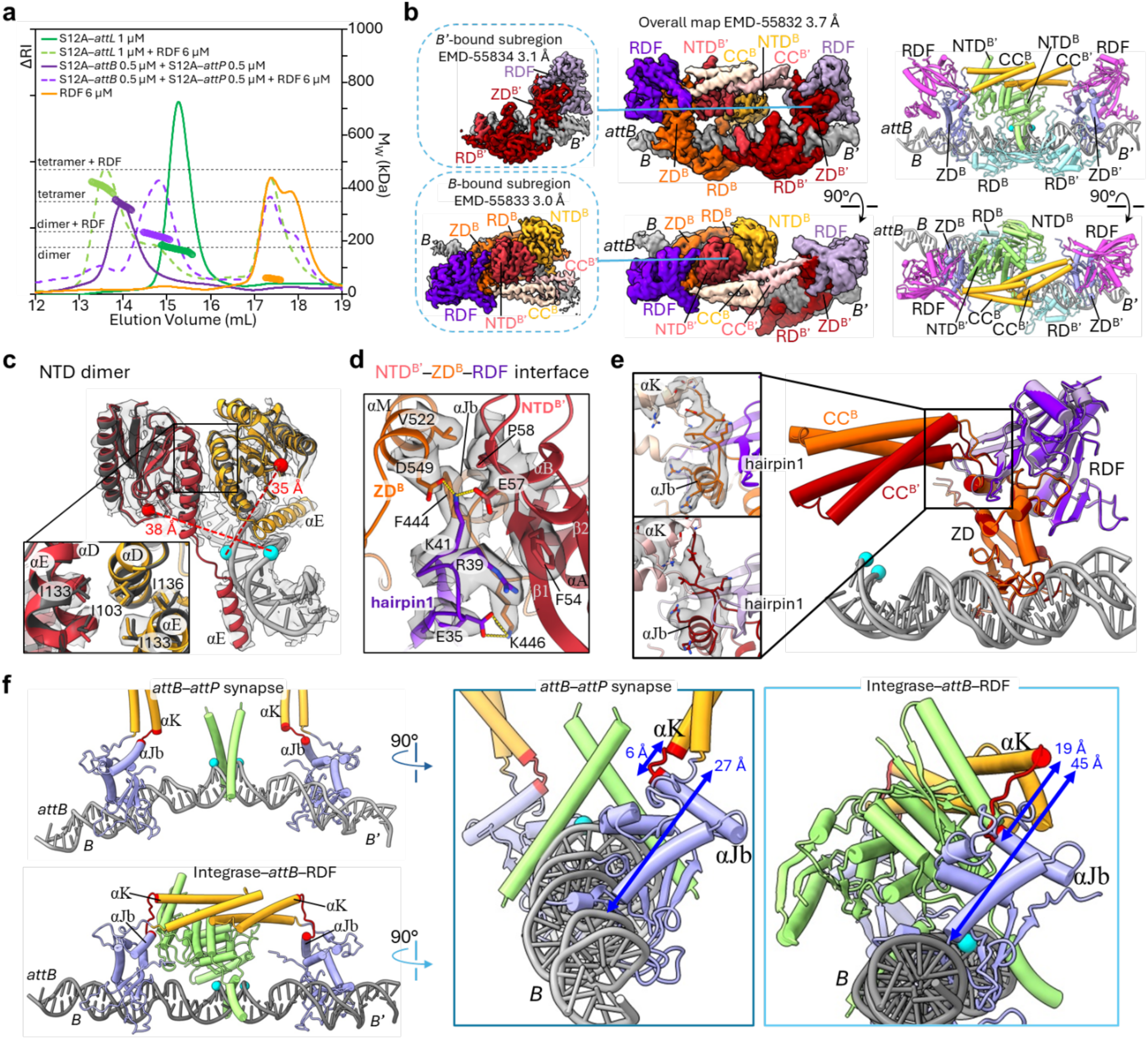
Structure of the integrase–*attB*–RDF dimer complex. **a**, SEC-MALLS analysis of the integrase^S12A^ complexes formed with *attL*, and with *attB* plus *attP*, in the absence and presence of excess RDF. Elution profiles are shown as solid or dashed coloured lines and M_W_ as circular markers. Dark green, *attL* without RDF; light green, *attL* with RDF, dark purple, *attP* and *attB* without RDF; light purple *attP* and *attB* with RDF; yellow, RDF by itself. **b**, Cryo-EM maps and the atomic model of the integrase^S12A^–*attB*–RDF complex. High-resolution local maps are in dashed boxes. **c**, Close-up view of the NTD dimer (red and yellow) and the central 16 bp of *attB* (light grey), with the cryo-EM maps overlaid in transparent grey. Cα atoms of the S12A residues are shown as red spheres, and the phosphorus atoms of the scissile phosphates as cyan spheres. Distances between the S12A residues and their respective scissile phosphates are indicated in red. For comparison, the NTD dimer of the integrase–*attL* complex (residues 1– 140) is superimposed in dark grey. The inset shows details at the dimer interface, with the side chains of I103 and I133 shown as sticks. **d**, Close-up of the NTD^B’^–ZD^B^–RDF interface. Side chains involved in the interface are shown as sticks, with their cryo-EM densities overlaid in transparent grey. **e**, Structural comparison of the CC motif trajectories of monomers bound at *B*- and *B’*. The panels on the left show details at the αJb–αK loop of each monomer, with their densities overlaid in transparent grey. The loop contacts hairpin1 of RDF, and undergoes lengthening upon RDF binding. **f**, Comparison of the structures of *attB*-bound monomers in the integrase^S12A^–*attB*–*attP* synaptic complex (left top) and the integrase^S12A^–*attB*–RDF complex (left bottom), with close-up orthogonal views shown on the right (boxed in dark and light blue respectively). For clarity, the integrase RD domains (residues 164–355) and the RDF proteins are hidden. The αJb–αK loops are highlighted in red. This loop is lengthened in the RDF-bound state and raises αK away from αJb and *attB* DNA. Distances from the N-terminus of αK to αJb and to DNA are indicated in blue.

We then solved the *attB*–integrase–RDF complex structure (Fig. 4b, Extended Data Fig. 8 and Methods). The structure reveals a centrally located intra-site CC dimer, positioned directly above the two scissile phosphates, approximately ∼29 Å away from the DNA (Fig. 4b). The NTDs form the same inactive dimer structure as was seen in the *attL*-bound dimer complex in the absence of RDF (Fig. 4c), and are asymmetrically packed against the ZD domain and RDF bound at the *B* half-site, creating a trimeric interface involving the *B’*-bound NTD and the *B*-bound ZD and RDF (Fig. 4d). The αJb–αK loop is expanded in a similar manner to that seen in the excisive synaptic complex (Fig. 4e), so that the start of αK is ∼45 Å from the *attB* DNA at both half-sites, compared with a much smaller distance of ∼27 Å in their RDF-free counterparts in the *attB*–*attP* complex (Fig. 4f). This RDF-mediated shift of the CC dimer away from the DNA avoids a steric clash with the paired NTDs, allowing the CC motifs to dimerise, locking *attB* in a dimeric, synapsis-deficient state. We speculate that a RDF-induced locked dimer also occurs for *attP*. In summary, our results illustrate how RDF-induced conformational changes of integrase activate *attL*/*attR* (excision) substrates by promoting synapsis, and inhibit integrative recombination by blocking *attB*–*attP* synapsis.

### Modelling of catalytic domain activation upon synapsis

Our structures of pre-cleavage synaptic complexes and inactive dimer complexes imply that synapsis and activation of integrase subunits for catalysis of DNA strand exchange require a dimer-to-tetramer transition involving remodelling of the integrase monomers and the dimer interface (Fig. 5a), analogous to the transitions seen in γδ and Sin serine recombinases^26^. To model the ready-to-cleave state, we generated an AlphaFold3^27^ (AF3)-predicted structure of the integrase NTD tetramer. The model has the expected 222 symmetry, and its αE arrangement closely matches that observed in our synaptic structures (Fig. 5b). The NTD core of each monomer in the AF3-predicted tetramer has a markedly different position from that observed in our integrase dimer–*att* site structures, rotated by ∼70° about αE toward its DNA-binding face (Fig. 5c). This arrangement positions the active site serine residue (S12) against the DNA-binding face of αE, similar to its placement in the activated γδ and Sin recombinases (Extended Data Fig. 9)^26^.

**Fig. 5:**
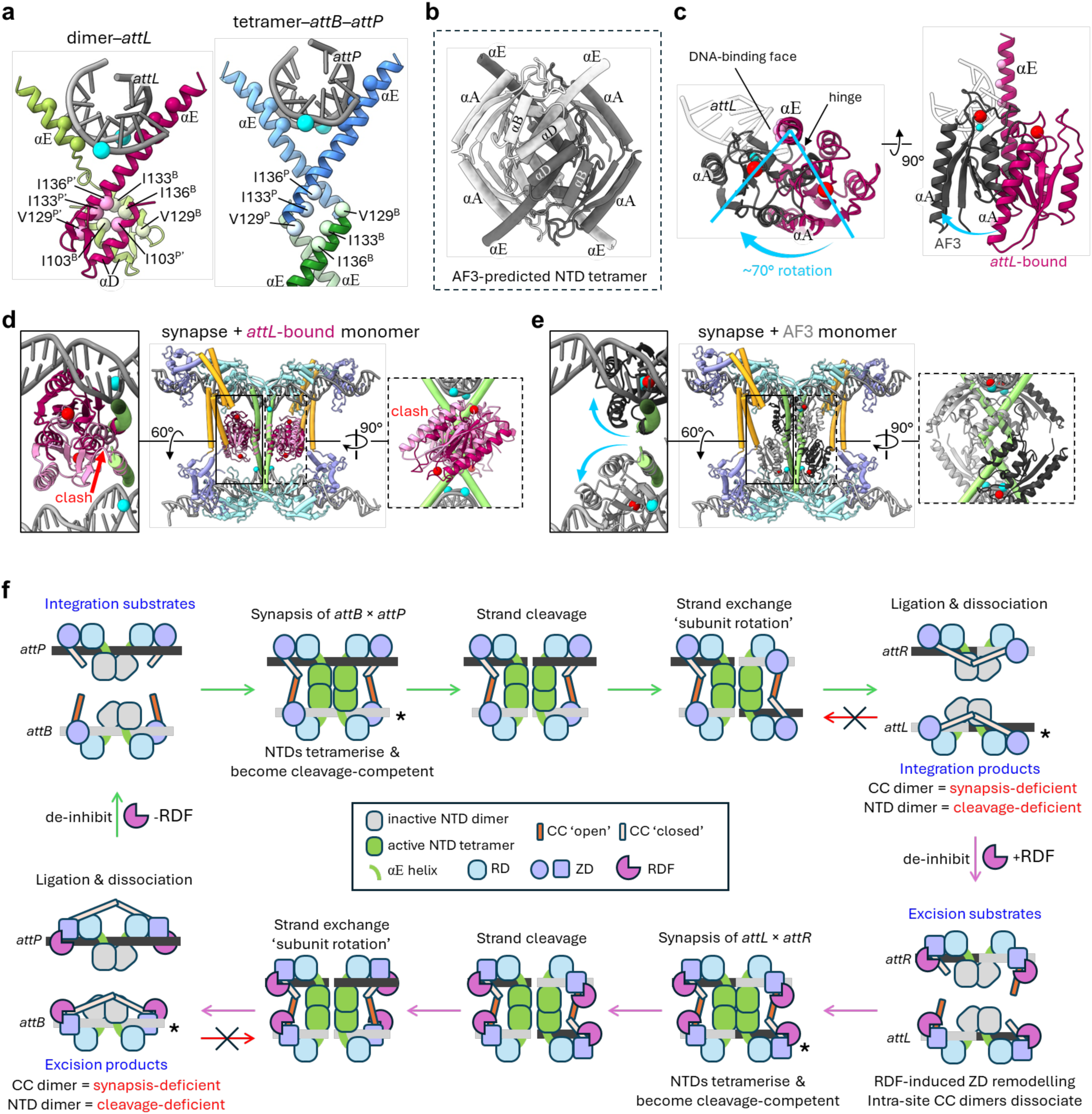
Catalytic domains undergo activating rearrangements upon synapsis. **a**, Structural comparison of the NTD dimer (left) and tetramer (right) interfaces, in the integrase–*attL* complex and integrase^S12A^–*attB*–*attP* synaptic complex respectively. For the tetramer, residues 129–163 (αE helix) are shown; for the dimer, residues 98–163 (αD, β4, β5, and αE) are shown. The Cα atoms of residues involved in each interface are shown as spheres, along with residues on the DNA-binding face of αE. Phosphorus atoms of the scissile phosphates are shown as cyan spheres. **b**, Tetrameric assembly of the full NTD domains (residues 1–163) predicted by AlphaFold3 (AF3) (ref. ^27^). A similar view to **a** is shown, with monomers in light and dark grey. **c**, Structural comparison of the *P’*-bound NTD monomer in the integrase–*attL* complex (magenta, residues 1–163) and the central 6 bp of *attL* (semi-transparent white) with a monomer in the AF3-predicted tetramer (grey). The structures are aligned at the N-terminal segment of αE (residues 129–140), with residues 141–163 of the AF3 monomer hidden for clarity. The DNA-binding residues of αE, the scissile phosphorus atom of the *P’* half-site, and the Cα atoms of S12 are shown as magenta, cyan, and red spheres, respectively. The blue arrow indicates a proposed ∼70° rotation of the NTD core around αE towards the DNA-binding face, a movement that aligns the catalytic serine with the DNA-binding face of αE where the scissile phosphate is positioned. The β5–αE loop that hinges the movement is indicated (black arrow). This region also facilitates an analogous conformational shift during the activation of γδ and Sin recombinases^26^. **d**,**e**, Modelling of the active NTD configuration within the synapse. The centre panels in **d** and **e** show the structure of the integrase^S12A^–*attB*–*attP* synaptic complex coloured by domain features as in Fig. 1b, overlaid with four copies of either the *P’*-bound NTD monomer of the integrase–*attL* complex (light and dark magenta, **d**) or the AF3-predicted monomer (light and dark grey, **e**). The structures are aligned at residues 129–140. The insets on the left and right show close-up views. The resulting model in **d** shows substantial steric clashes of the integrase monomers (red arrow), and the S12 Cα atoms (red spheres) positioned 17–22 Å from the scissile phosphates (cyan spheres). In contrast, the model in **e** shows near-complete resolution of the steric clashes, and places all four S12 residues adjacent to their respective scissile phosphates (<8 Å, Cα–P). **f**, Structure-based model of unidirectional recombination by serine integrases, showing the four key reaction steps — synapsis, strand cleavage, strand exchange, and ligation — during both integration and excision pathways. Cartoons illustrate various integrase–*att* and integrase–*att*–RDF complexes formed during the recombination cycle, with those structures reported in this study marked with an asterisk. See Discussion for details.

**Fig 6:**
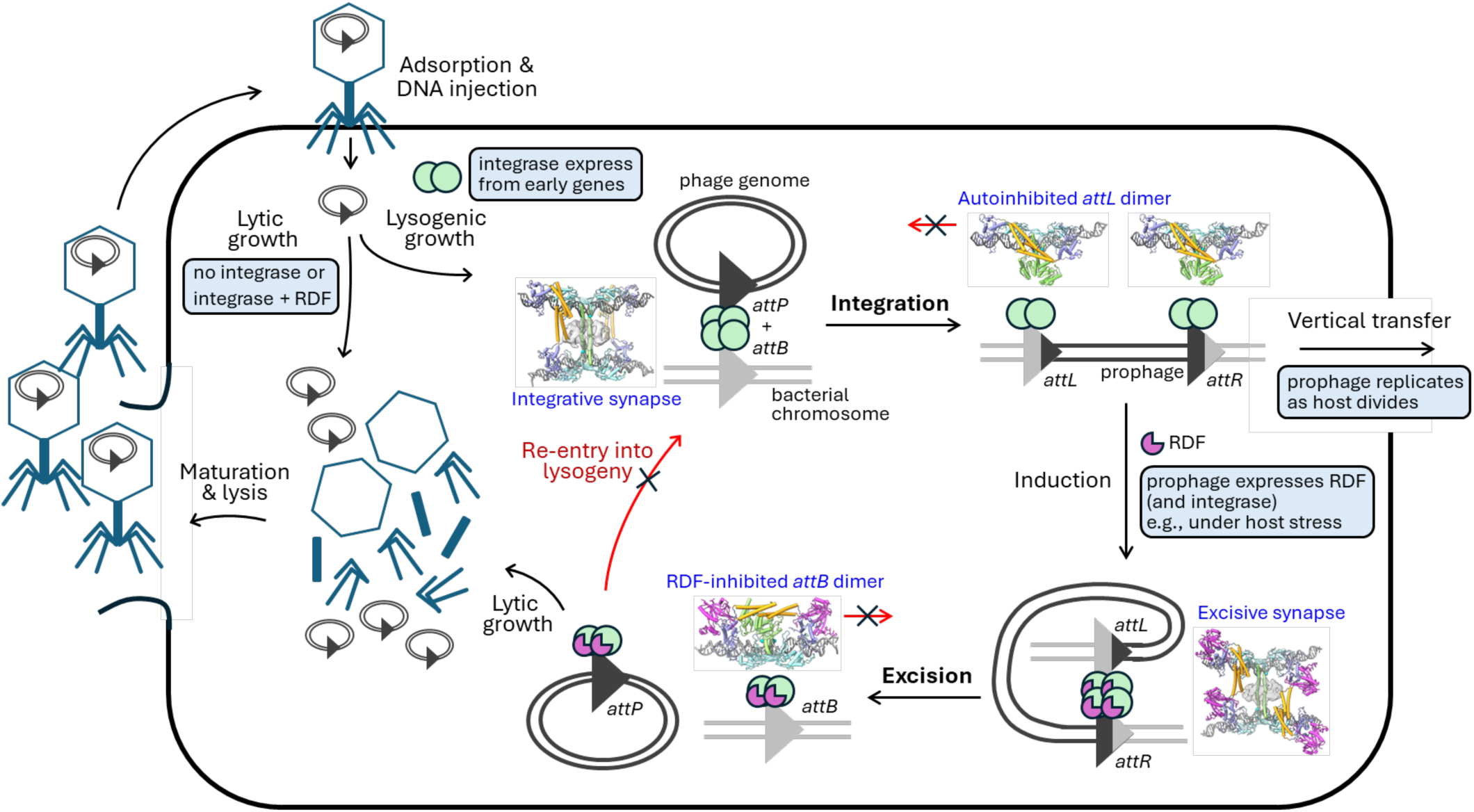
Model of serine integrase-mediated recombination in bacteriophage lysogeny. Temperate bacteriophage (phage) infections begin with adsorption of the virus to the host cell and injection of the viral genome into the cytoplasm. Here the virus makes a decision to either replicate *via* the lytic cycle, where host replication machinery is hijacked to make new viral particles, or enter the lysogenic cycle, with the virus existing as a stably integrated prophage that replicates as the host divides. In phages with a serine integrase system, expression of integrase drives entry into lysogeny and, given that RDF inhibits *attB* × *attP* recombination, we reason that commitement to lytic growth can arise either from repression of integrase or expression of RDF. Lysogenic prophages can then replicate passively as their hosts divide. Induction occurs when integrase and RDF are both expressed, typically in response to host stress, leading to excision of the prophage. The excised phage is prevented from re-integrating into the chromosome through RDF-mediated inhibition of *attP* × *attB* recombination, thereby commiting the virus to lytic replication.

To visualise the dimer-to-tetramer transition, we first placed a cleavage-deficient integrase monomer (bound to the *P’* half-site of *attL*) into the *attB*–*attP* synaptic assembly by aligning residues 129–140 from the N-terminal portion of αE. This produced pronounced steric clashes between the NTD cores within each rotating dimer (Fig. 5d). Repeating the same procedure with the AF3-predicted monomer instead showed that the ∼70° NTD rotation resolves these clashes and brings the active site serine residues to within ∼8 Å of their respective scissile phosphates (Fig. 5e). This model therefore likely represents the ready-to-cleave state of the synaptic complex.

## Discussion

Here, we present the first complete structures of intermediates in serine integrase-mediated recombination. Our four structures represent key stages in catalysis and regulation, and elucidate the mechanism by which unidirectional recombination is achieved (Fig. 5f).

As previously hypothesized (refs. ^13-16^), the crucial checkpoint is at synapsis of two *att* sites. In both of our synaptic complex structures, the NTDs (catalytic domains) of the integrase subunits bound at the two *att* sites interact to form a tetramer at the site centres, close to the DNA phosphodiesters that are to be cleaved and rejoined in recombination. For small serine recombinases such as γδ resolvase, the NTD tetramer is sufficient to maintain synapsis of the two sites, but this is not the case for integrase, which requires additional bridging (inter-site) CC–CC interactions (Fig. 1d,i). These bridging interactions must be commensurate with a distance between the two *att* sites at their centres (∼50 Å) which allows engagement of the NTD active sites with their target phosphodiesters. The ’hinge region’ that links the CC motif with the main body of the ZD (of which it is a part) is very flexible, allowing it to adopt a wide range of angular trajectories (Fig. 1k). Furthermore, the α-helical secondary structure of the CC motif adjacent to the hinge can partially disassemble (Fig. 2d), resulting in extension of the motif and thus allowing contacts between the tips of pairs of CC motifs whose anchor points are further apart (Figs. 1d and 2b). Due to this structural plasticity the CC motifs can make interactions between subunits bound at *attB* and *attP* sites leading to a productive synaptic complex (Fig. 1d), but productive interactions are not achievable for other combinations of *att* sites. However, a productive synaptic complex can be formed between *attL* and *attR* sites in the presence of the RDF, which binds to the ZD hinge region of each integrase subunit to change the position, length and trajectory of the CC motif (Fig. 3 and Supplementary Video 5). Importantly, the sites are aligned in this synaptic complex to yield *attB* and *attP* recombination products, and thus ensure phage excision.

The CC motifs of the two subunits bound to a single *att* site can interact if the separation (taking account of DNA bending) and helical phase of the *att* site-bound ZDs permit it, and there is not steric hindrance by the DNA or other parts of the integrase subunits, as shown in our *attL*– integrase dimer structure (Fig. 2b). The intra-site CC–CC interaction renders the motifs unavailable for inter-site synaptic interactions and thus inhibits recombination. We presume that inactive *attL* and *attR* complexes form rapidly upon completion of *attB* × *attP* recombination, on dissociation of the integrase-bound product sites. In contrast to the *attL* complex, the CC motifs of an integrase dimer bound to *attB* apparently do not make a stable interaction in the absence of RDF. However, the structural changes induced by RDF binding to the integrase hinge regions allow the motifs to escape from the steric hindrance of the NTD dimer at the centre of the site and make a stable interaction (Fig. 5f). Thus RDF functions both to promote *attL* × *attR* recombination by enabling synapsis, and to inhibit *attB* × *attP* recombination by rendering the *attB*-integrase dimer synapsis-deficient. We speculate that RDF might also promote intra-site CC–CC interaction in *attP*-bound integrase dimers by altering the integrase conformation at the base of the CC motifs in an analogous way. Interestingly, although the two RDFs bind to similar integrase regions, the specifics of the ϕC31 integrase:RDF interaction appear different from that of SPbeta integrase^28^.

In addition to the inhibitory intra-site CC–CC interaction, our two dimer-bound structures also reveal an interface comprising the αD and αE helices that dimerises the NTDs into a cleavage-deficient conformation (Figs. 2h and 4c), potentially providing a further block to any catalytic (DNA cleavage) activity. The NTD dimer interface engages surfaces that overlap with those used in the tetramer assembly within the synaptic complex (Fig. 5a), suggesting that synapsis is required to dissociate the inactive dimer and induce an active conformation. Analogous inactive NTD dimers have been observed in structures of the small serine recombinases γδ resolvase and Sin (ref. ^26^).

In summary, recombination between *att* sites is dependent on synaptic complex formation stabilized by inter-site CC–CC interactions, which in the absence of RDF are favourable only for *attB*–*attP* site pairs, but in the presence of RDF are favourable only for *attL*–*attR* pairs. For each reaction, recombination of the product *att* sites is inhibited, thus implementing unidirectionality.

We anticipate that the data presented here will pave the way towards structure-guided engineering of serine integrases for DNA manipulation applications, including those that exploit the previously under-utilized switching of recombination directionality by RDF proteins.

## Acknowledgements

We acknowledge support from BBSRC grant BB/W017571/1 to LS, MWS and SDC and of a College of Medicine Veterinary Medicine and Life Sciences PhD studentship to L.S. Preliminary data were collected at the Scottish Centre for Macromolecular Imaging (SCMI), University of Glasgow, funded by the MRC (MC_PC_17135, MC_UU_00034/7, MR/X011879/1) and SFC (H17007) and supported by James Streetley. We acknowledge Diamond Light Source for access to the cryo-EM facilities at the UK national electron Bio-Imaging Centre (eBIC), proposal BI37630, supported by Parijat Majumder, Kyle Dent, Cecile Fort, Yakun Tang and Dr Dan Clare. We acknowledge the MVLS Advanced Research System (MARS) at the University of Glasgow for providing high-performance computing resources and technical support. We are particularly grateful to Jasmin Schar (MARS). We acknowledge Michal Klosowski and Gemma Harris (Research Complex at Harwell) for technical assistance with cryo-EM and SEC-MALLS. We acknowledge experimental support from Andrew Gunathilaka (Imperial College London) and Anna Cameron (University of Glasgow).

## Contributions

Y.E.S. designed the expression constructs for unpublished plasmids, purified the protein samples, assembled the protein:DNA assemblies, prepared cryo-EM grids, collected cryo-EM data, calculated cryo-EM maps, built and refined structural models, performed SEC-MALLS experiments, prepared figures and wrote the initial article draft. L.A. conducted preliminary sample preparation and performed preliminary single-particle cryo-EM experiments. A.P.J. performed model building and refinement. L.S. performed preliminary cryo-EM data collection and analysis. L.S., W.M.S. and S.D.C. conceptualized and oversaw the project, obtained funding and analysed data along with the other authors. All authors contributed to the drafting and revision of the manuscript.

## Methods

### Protein expression and purification

Proteins used in this study, including ϕC31 integrase (Uniprot ID: Q9T221), integrase variants (S12A, S12A/Y475H, S12A/D477K, S12A/E449K, ΔNTD, S12A/ΔCC), the RDF gp3 protein (Uniprot ID: Q9T216), and the integrase^S12A^-RDF fusion construct, were expressed in the *Escherichia coli* BL21 (DE3) LOBSTR^29^ strain using pET28a expression vectors with synthetic codon-optimised coding sequences. All constructs contained an N-terminal hexahistidine purification tag. The integrase^S12A^-RDF fusion construct connected the N-terminus of RDF to the C-terminus of integrase with a 32-residue GS-rich flexible linker (*VAA-*TSGTSGGGTGAGSGTGSGGSGSGTAGGSTAGG-*MAK*). A single transformed colony was used to inoculate an overnight pre-culture grown in LB media at 37 °C, shaken at 180 rpm. Pre-culture was used to seed 1 L LB medium cultures at a 1:200 ratio, which were grown at 37 °C, shaken at 180 rpm. Protein expression was initiated by the addition of 0.5 mM IPTG to log phase cultures prior to subsequent growth overnight at 18 °C, 180 rpm. Cultures were harvested by centrifugation the next morning, washed once with 20 mM HEPES, pH 7.4, 150 mM NaCl, and stored at -20 °C until purification.

To purify, cells were resuspended 1:5 (w/v) in lysis buffer (20 mM HEPES, pH 7.4, 1 M NaCl, 2 mM MgCl_2_, 10 mM imidazole, 10% (v/v) glycerol, 0.1% (v/v) Tween 20, 0.5 mM TCEP, 1 × cOmplete^TM^ mini protease inhibitor tablet (Sigma), 1 × benzonase (Sigma)), then lysed by sonication, and centrifuged (Sorvall F21-8×50y, at 4 °C, 20,000 rpm, 30 min). Clarified lysate was then passed through a 1 mL HisTrap Ni-NTA column (Cytiva) to capture the His-tagged proteins. The bound proteins were washed with 30 column volumes (CV) of IMAC buffer A (20 mM HEPES, pH 7.4, 1 M NaCl, 2 mM MgCl_2_, 10 mM imidazole, 0.5 mM TCEP), and subsequently eluted with an imidazole gradient of 10–300 mM over 10 CV. Protein-containing fractions were analysed by SDS-PAGE, and those with high purity were concentrated in a Amicon® spin concentrator (10 or 50 kDa MWCO). Concentrated protein samples were further purified by size-exclusion chromatography. For integrase, integrase variants, and the integrase^S12A^-RDF fusion, a Superose 6 increase 10/300 column was used, equilibrated in 20 mM HEPES, 1 M NaCl, 2 mM MgCl_2_ and 0.1 mM TCEP (pH 7.4). For RDF, a Superdex 200 increase 10/300 column was used, equilibrated in 20 mM HEPES, 150 mM NaCl, 2 mM MgCl_2_ and 0.1 mM TCEP. Eluted protein fractions were analysed by SDS-PAGE, and those with high purity were pooled and concentrated to 10–30 mg/ml, flash frozen and stored at -80 °C.

### Assembly and purification of integrase–att complexes

Integrase–*attB*, integrase–*attP*, and integrase–*attL* complexes were freshly assembled before SEC-MALLS and cryo-EM experiments. To assemble the complexes, the *att* site duplex was mixed with an equimolar amount of integrase, with a final buffer composition of 20 mM HEPES, 150 mM NaCl, 2 mM MgCl_2_ and 0.1 mM TCEP (pH 7.4). The mixture was incubated at 30 °C for 15 min to allow complex formation, then centrifuged at 15,000 rpm for 10 min at 4 °C to remove precipitates before injection onto a Superose 6 column equilibrated in the same buffer. Fractions that contained pure complexes were identified by SDS-PAGE. Given that the DNA component contributes the vast majority of the absorbance at 260 nm, the molar concentration of the integrase–*att* complex was taken as the *att* site DNA concentration, determined from A_260_ measurements. The same steps were taken to assemble and purify the excisive *attL*–*attL* synaptic complex using *attL* DNA with the integrase^S12A^-RDF fusion protein.

The full *attB*–*attP* integrative synaptic complex was made by mixing equal molar amounts of purified integrase–*attB* and integrase-*attP* complexes, and incubating the mixture at room temperature for 5 min prior to SEC-MALLS injection or cryo-EM vitrification. The RDF-bound complexes were prepared by mixing purified integrase–*att* complexes, or the *attB*–*attP* synaptic complex, with RDF at 1:3 integrase:RDF molar ratio, followed by a 5-min incubation at room temperature before SEC-MALLS injection. The integrase–*attB*–RDF complex used in cryo-EM was further purified by SEC to remove excess RDF.

### SEC-MALLS

SEC-MALLS was used to determine the solution stoichiometry of integrase–*att* and integrase–*att*– RDF complexes. Samples (100 μL) were applied to a Superose 6 increase 10/300 column mounted on an AktaPure 25 (Cytiva), equilibrated in 20 mM HEPES, 150 mM NaCl, 2 mM MgCl_2_ and 0.1 mM TCEP (pH 7.4) at a flow rate of 0.5 mL/min and a temperature of 25 °C. The scattered light intensity and macromolecular complex concentration of the column eluate were recorded using a Wyatt Dawn HELEOS-II 18-angle light scattering detector and a Wyatt Optilab T-rEX differential refractive index detector (ΔRI) (dn/dc = 0.180), respectively. The weight-averaged molecular mass (M_W_) of material contained in chromatographic peaks was determined from the combined data from both detectors using the software ASTRA (Wyatt Technologies).

### Cryo-EM sample preparation

Cryo-EM grids were prepared using purified complexes, including the *attB**–**attP* synaptic complex (0.25 mg/ml), integrase***–****attL* complex (0.4 mg/ml), *attL**–**attL* synaptic complex (0.5 mg/ml), and integrase***–****attB**–***RDF complex (0.4 mg/ml). All complexes were prepared using the cleavage-deficient S12A integrase variant, except for the integrase***–****attL* complex where the WT integrase was used. Complexes were prepared fresh before vitrification as described above. Samples (4 μL) were applied onto a freshly glow-discharged Quantifoil R 1.2/1.3 copper 300 mesh grid. The sample was then blotted for 2 s at a blot force of 6 and vitrified by plunging into liquid ethane using a Vitrobot Mark IV (FEI Company) operated at 4 °C and 100% humidity.

### Cryo-EM data collection

All cryo-EM data sets were collected on FEI Titan Krios transmission electron microscopes operated at 300 keV at the UK national cryo-EM centre eBIC. Multi-frame movie stacks were collected at a total electron dose of 50–60 e^-^/Å² and a dose-per-fraction of ∼1 e^-^/Å², yielding data sets containing 7,611 to 27,020 movies. A nominal pixel size of 0.504 to 1.171 Å was used, with a defocus range of-0.7 to -2.5 μm. Data set-specific collection optics and image processing parameters are summarised in Extended Data Tables 1 and 2.

### Cryo-EM image processing

All data sets were processed in RELION^30^ v.5.0 with BLUSH regularisation^31^ used in image alignments. Motion correction was carried out using RELION’s implementation, and the resulting micrographs were contrast transfer function (CTF) corrected using CTFFIND-4.1^32^. Both global and local reconstruction schemes were applied to each data set, yielding global reconstructions at 7.2 Å, 4.4 Å, 7.2 Å, and 3.7 Å resolution, and local reconstructions at 3.2–3.8 Å, 3.4–3.9 Å, 3.1–3.3 Å, and 3.0–3.1 Å resolution, for the *attB*–*attP* synaptic complex, *attL*–integrase complex, *attL*–*attL* synaptic complex, and integrase–*attB*–RDF complex, respectively. All resolutions are reported based on the gold-standard FSC = 0.143 criterion^33^ estimated from RELION’s postprocessing job. CTF refinement and Bayesian polishing were performed on all final particle stacks for reconstructions that reached a resolution of 4.4 Å or better.

### attB–attP synaptic complex data set

Particles were picked from all CTF-corrected micrographs with RELION’s LoG-based autopick programme, which yielded an *ab-initio* 3D map after a round of classification and selecting those that generated good 2D averages. Particle picking was then repeated with RELION’s template-matching algorithm using the *ab-initio* map as a 3D reference model, yielding 1,921,071 picked coordinates from 13,330 micrographs. Particles were extracted with a binned pixel size of 1.5 Å and a box size of 128. Four successive rounds of 3D classification with a global mask and the selection of the most well-assembled classes yielded a final particle stack of 11,085 particles. The unbinned particle images (1.171 Å/pix, box size 320) were then used to obtain a final map of the fully assembled *attB*– *attP* synaptic complex at 7.2 Å resolution.

Local classification and refinement^17^ were employed to obtain high-resolution reconstructions of the *attB*-and *attP*-bound subregions. First, template-based particle picking was repeated with the 7.2-Å global map as the reference and a more generous picking threshold, yielding 6,090,789 particles. Particles were subjected to one round of global 3D classification, yielding 1,950,242 non-junk particles. Subsequent rounds of 3D classification with local masks covering the individual *attB*-or *attP*-bound regions yielded final particle stacks containing 274,520 and 253,292 particles, respectively, with each refining to 3.2 Å and 3.3 Å resolution. In some cases, signal subtraction^17^ of the unmasked regions was employed to further improve classification efficacy, but particles were always reverted back prior to refinement. The detailed workflows used are summarised in Extended Data Figure 1.

A third 3.8-Å local reconstruction was obtained that showed well-defined details at the CC– CC dimer interface. This was obtained by applying successive *attB*-and *attP*-bound local masks on the 1,950,242 non-junk particles to yield 226,040 particles that showed strong densities at both regions. These were further classified using a mask that covered a single *P’*-bound monomer and the CC motif from the opposing *B’*-bound monomer. The final particle stack contained 29,776 particles that reconstructed to 3.8 Å.

### Integrase–attL complex data set

Similar steps were taken to generate an *ab-initio* map, except that picking was done on a subset of 1,000 micrographs using RELION’s built-in Topaz^34^ picker (untrained). The particle stack was cleaned through two rounds of global classification, yielding a 7.1 Å reconstruction from 11,846 particles. These were used as the 3D reference and Topaz training particle set to pick from all 8,830 micrographs, yielding 5,885,905 and 5,984,437 particles from the reference-based and Topaz (trained) pickers, respectively. Classification was performed on each particle stack, first with a global mask, then with a local mask that contained the *attL* duplex and the DNA-bound domains. The selected particles were combined and 115,312 duplicate picks were removed using a 30-Å distance threshold, yielding 713,127 particles that reconstructed to 3.8 Å. Global and local masks were then applied for further 3D classification, yielding a global reconstruction at 4.4 Å from 14,425 particles. Two locally refined maps at 3.4 Å and 3.9 Å were also obtained for the *B*-and *P’*-bound subregions, from 122,514 and 35,608 particles, respectively. All final reconstructions were generated using unbinned particle images with a pixel size of 1.072 Å and a box size of 256. Detailed processing workflows are summarised in Extended Data Figure 4.

### attL–attL synaptic complex data set

Similar procedures were used to generate an *ab-initio* map using Topaz-picked (untrained) particles from all 7,611 micrographs. The particles were further classified to yield a 8.5 Å reconstruction of the full complex from 8,332 particles, which were used as inputs to repick 2,430,741 and 1,928,306 particles using reference-based and Topaz (trained) pickers, respectively. Particles of the two picking sets were extracted with a binned pixel size of 5.8 Å and a box size of 64, and each underwent two rounds of global 3D classification. One class from each picking set was selected and combined for further global analysis, whereas for the local reconstructions, a more permissive selection of three classes per picking set was applied to retain those with good local but less complete global densities. Duplicate picks were removed from each combined set (<30 Å threshold), yielding 410,587 and 961,367 for the global and local sets, respectively (Extended Data Fig 3). For the global set, four subsequent rounds of 3D classification yielded a final reconstruction of the full complex at 7.2 Å from 17,432 particles. For the local set, particles were aligned to their C2 symmetry axis and symmetry expanded with C2 symmetry, yielding 1,922,734 particles in the expanded set. Local masks were applied to a single *B*- and *P’*-bound monomer, and signal subtraction was carried out to remove particle regions outside the masks. The subtracted subregions were subjected to two rounds of local 3D classification, yielding 241,788 and 118,011 particles for the *B*- and *P’*-bound subregions, respectively. To avoid cases in which a symmetry-related partner reverts orientation during classification or refinement, all symmetry-related duplicates were removed (<30 Å threshold), retaining only one particle from each pair. The resulting particle stacks underwent one additional round of 3D classification, yielding final stacks of 115,490 and 66,541 particles for the *B*- and *P’*-bound subregions, respectively, and producing final reconstructions at 3.1 and 3.3 Å. All final reconstructions were generated using unbinned particle images with a pixel size of 1.06 Å and a box size of 360. Detailed processing workflows are summarised in Extended Data Figure 6.

### Integrase–attB–RDF complex data set

An *ab-initio* model was generated with untrained Topaz picked particles from a subset of 1,000 micrographs. Particles then underwent two rounds of 3D classification to produce a 6.6-Å map of the full complex from 13,874 particles, which were then used as inputs to repick 2,727,541 and 5,833,916 particles using reference-based and trained Topaz pickers, respectively. Each picking set underwent two rounds of global 3D classification, producing a combined count of 521,264 good particles after removing 44,852 duplicate picks (<30 Å threshold). Global and local masks were then applied, producing a 3.7-Å map of the full complex from 30,771 particles, and a 3.0-Å and 3.1-A map for the *B*- and *B’*-bound subregions, respectively, from 60,721 and 87,707 particles. Final reconstructions were generated with a pixel size of 1.008 Å and a box size of 280. Detailed processing workflows are summarised in Extended Data Figure 8.

### Model building and refinement

Prior to this study, the only experimental structure available for ϕC31 integrase was a crystal structure of the NTD-RD fragment (PDB 4BQQ). We therefore generated AF3-predicted models of full integrase dimers bound to *attB* and *attP* as starting models. Both models showed good agreement with known integrase domain structures. Additionally, the *attP* model correctly reproduced the domain-DNA positioning seen in LI integrase bound to an *attP* half-site. However, the 5-bp binding-register shift of the *attB* ZD was not correctly predicted. The two starting models were then manually fitted into their respective 3.2-Å and 3.3-Å local maps (EMD-52736 and EMD-52737) in ChimeraX^35^ and Coot^36^. Model regions that lacked density were removed, including the NTD residues 1–139 and residues 590–605 at the C-terminus. Additionally, residues 450–518 (CC motif) were removed for the *attB*-bound monomers, and base-pairs -/+28–30 removed for *attP*60 DNA. In both maps, the map quality was sufficient to unambiguously assign the orientation of the *attB53* and *attP60* substrates. The models were then iteratively refined through manual adjustment in Coot followed by automated refinement in Phenix^37^ with secondary-structure restraints, until satisfactory MolProbity^38^ validation statistics were achieved. The refined *attB*- and *attP*-bound models were deposited into the PDB with accession codes 9I8S and 9I8T, respectively. These were used as starting models to refine all other integrase–*att* structures.

To refine the full *attB*–*attP* synaptic complex, starting models 9I8S and 9I8T were fitted into the 7.2-Å global map (EMD-52811) using TEMPy-ReFF^39^. Models for the two *attB*-bound CC motifs were docked with EM-placement in the CCP-EM^40^ package, using the structure of the *attP*-bound CC motif as the starting model. The four αE helices were extended from residue D140 to their N-terminal residue V129, using the same region in 4BQQ as a starting model. Automatic model refinement was then performed in Refmac-Servalcat (CCP-EM) using external restraints generated from the starting models with ProSMART^41^ and LibG^42^. We could assign the orientation of the *attB* DNA, as the *attB53* substrate used was 3 bp longer on its left half-site (the *B* half-site) than on its right half-site (*B′* half-site). However, the map resolution was insufficient for unambiguous determination of the *attP60* DNA orientation. We modelled it in a ’parallel’ orientation relative to *attB*, as predicted for a synaptic complex that can lead to recombination.

Similar procedures were used for the remaining three complexes: models were first refined into their respective local maps, followed by refinement into the global map under strict model restraints. For the NTD dimer in the integrase–*attL* and integrase–*attB*–RDF complexes, the starting model was 4BQQ, whereas the starting model for the RDF (gp3) was generated using AF3. Model refinement and validation statistics are summarised in Extended Data Tables 1 and 2.

---

**Extended Data Fig. 1:**
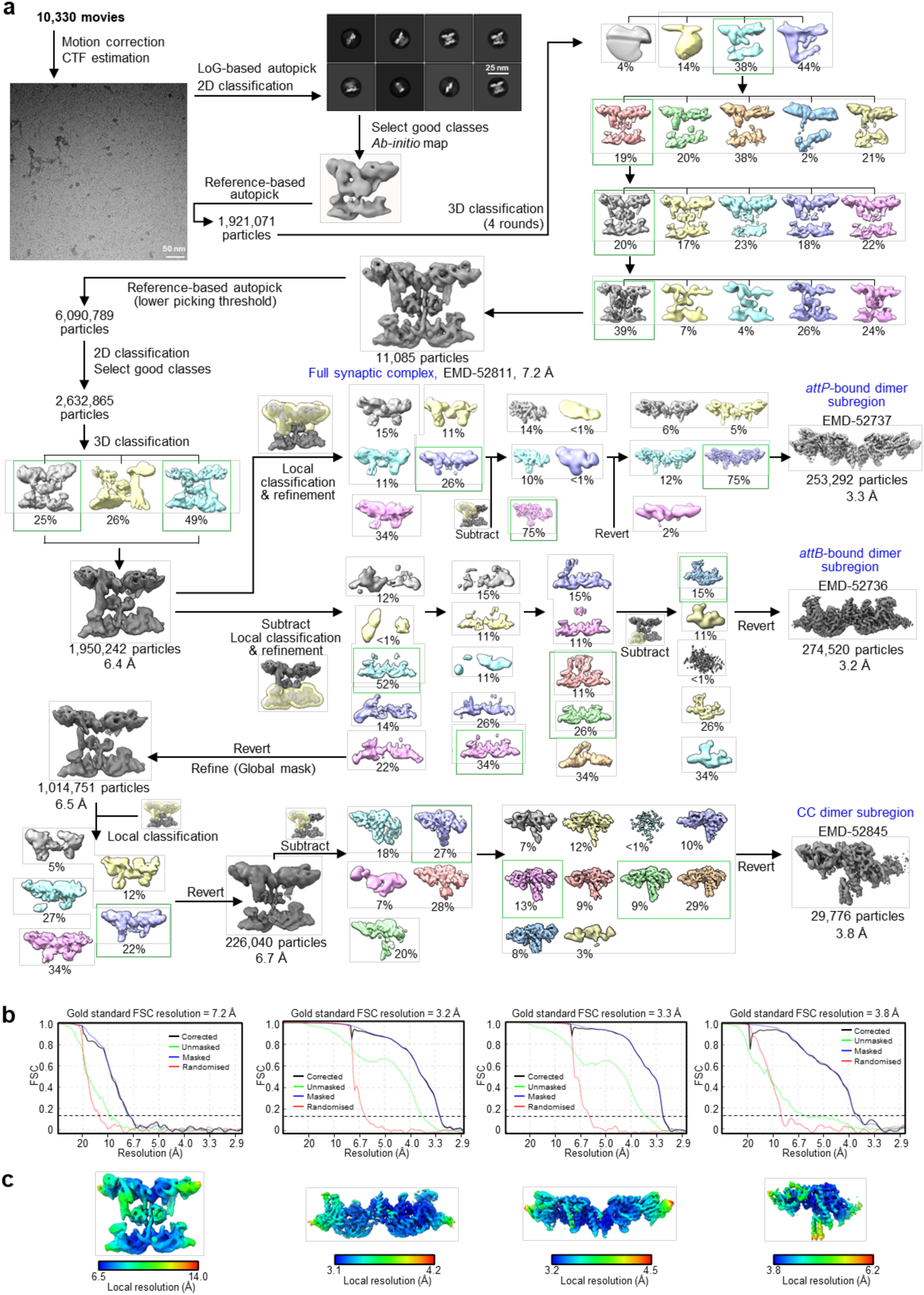
CryoEM analysis workflow and map quality metrics of the ϕC31 integrase^S12A^– *attB*–*attP* synaptic complex. **a**, Cryo-EM image processing workflow. Particle classes selected for further processing are boxed in green, and final maps used for model building are labelled in blue. Masks used for local reconstruction, classification, or signal subtraction are shown as transparent yellow overlays. **b**,**c**, Gold-standard FCS curves used for global map resolution estimation (**b**) and local resolution estimate (**c**) of the final maps. Left to right: full synaptic complex; *attB*-bound dimer subregion; *attP*-bound dimer subregion; *attP*-bound monomer showing well resolved CC-CC dimer.

**Extended Data Fig. 2:**
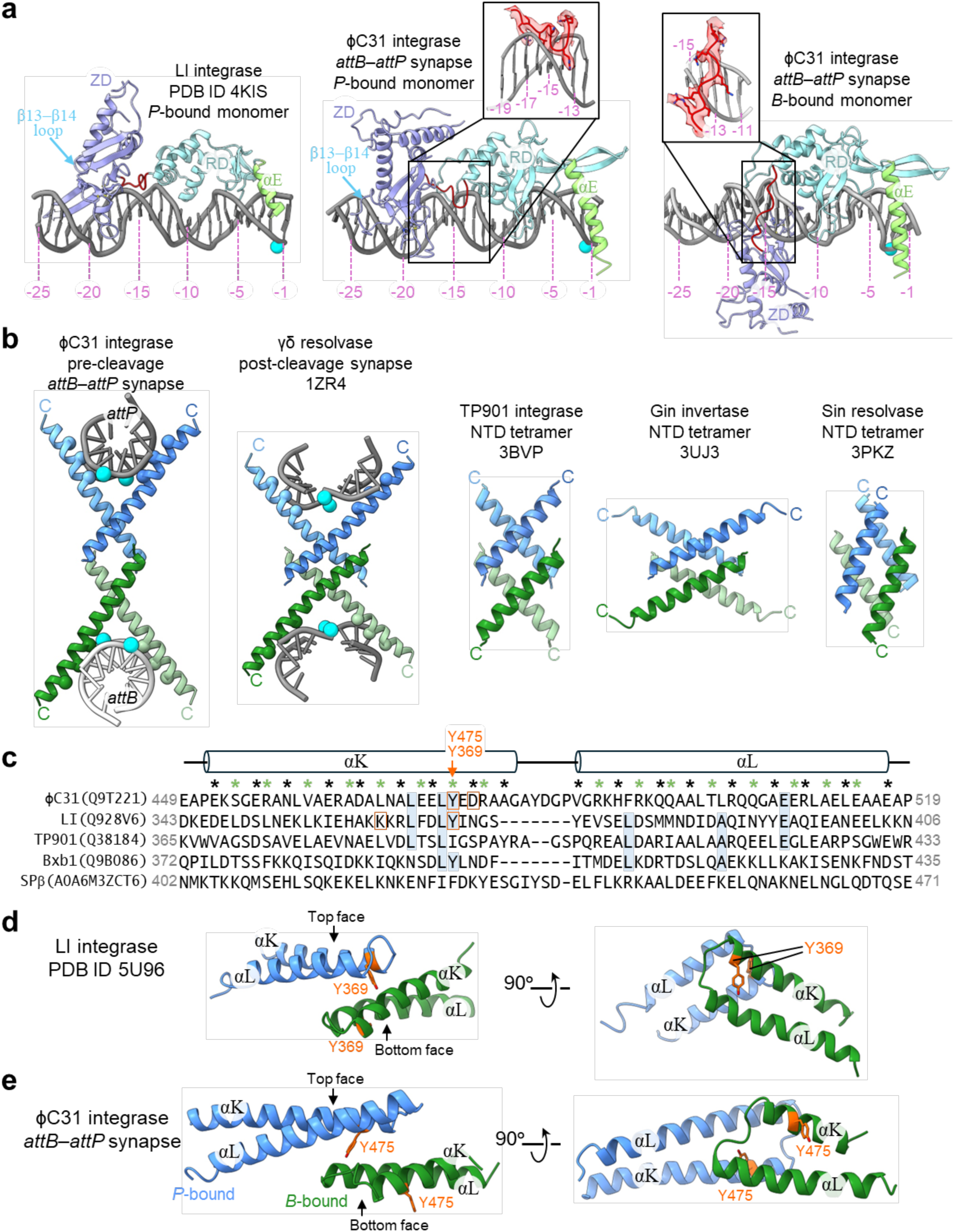
Structural comparisons of ϕC31 integrase with other serine recombinases. **a**, Structure of LI integrase bound to *attP* half-site (left, PDB 4KIS) compared to *P*- and *B*-bound monomers in the ϕC31 *attB*×*attP* synapse (middle and right). DNA is shown in grey with scissile phoshourous as cyan spheres. Protein regions are coloured as in Fig 1b, except the RD–ZD linker (residues 345–354) is highlighted in red. For clarity, CC motifs are hidden. Base-pair positions have been indicated in pink. Insets show close-ups of the RD–ZD linkers, with their cryo-EM density overlaid in transparent red. Blue arrows mark the β13–β14 loop that makes DNA contact in ϕC31 but not in LI. **b**, Comparison of the αE tetramer in serine recombinase structures; left to right: ϕC31 *attB*×*attP* synapse, γδ resolvase post-cleavage synapse (1ZR4), Gin invertase NTD tetramer (3UJ3), TP901 integrase NTD tetramer (3BVP), and Sin resolvase NTD tetramer (3PKZ). C-termini are indicated with letter C. For the synapse structures, the central eight nucleotides are shown as grey sticks, the Cα of DNA-binding residues as green and blue spheres, and the phosphorous atoms of the scissile phosphates as cyan spheres. **c**, Sequence alignment of the CC region from selected serine integrases, with the corresponding structure from ϕC31 shown above the alignment. Protein names are shown on the left, with their Uniprot IDs in parentheses. Amino acid boundaries are indicated at the sequence ends. Identical positions in three or more sequences are highlighted in blue. Residues on the ‘top’ and ‘bottom’ faces of CC, as observed in ϕC31 and LI structures, are marked with black and green asterisks, respectively. Orange boxes mark residues that disrupt CC–CC dimer in ϕC31 and LI^11^ when mutated. **d**,**e**, Structural comparison of the CC dimer in LI (**d**) and ϕC31 (**e**), with the top and bottom faces of CC indicated with black arrows. The bottom face residues, Y369 in LI and Y475 in ϕC31, are shown as orange sticks.

**Extended Data Fig. 3:**
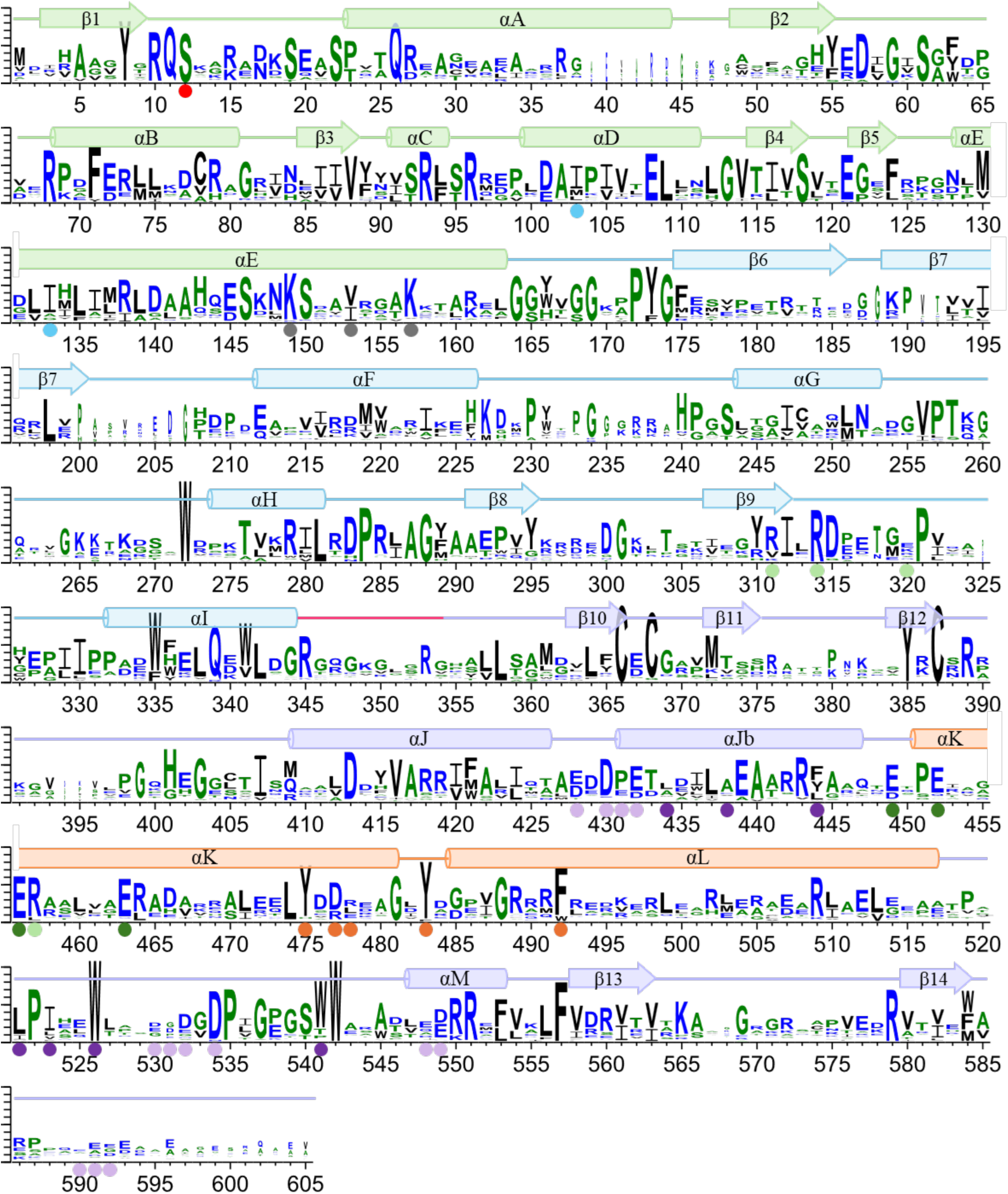
Sequence conservation of serine integrases in the *Streptomyces* genus. Sequence logos in one-letter code displaying amino acid conservation generated by WebLogo^43^, showing the results of an alignment of 46 serine integrase sequences obtained from *Streptomyces* genomes. Amino acid positions on the y-axis are numbered according to ϕC31, with their secondary structures shown above (α-helices as cylinders and β-strands as arrows). The height of the logos correspond to the degree of conservation at that position among the 46 sequences. Sequence were obtained by an initial BLAST^44^ search with ϕC31 integrase as the query against the NCBI RefSeq Select^45^ database, with results limited to 100 *Streptomyces* sequences. A filter was then applied to only include sequences with 30–60% identity to the query and 560–620 amino acids in size. Finally, sequences were inspected to ensure the presence of complete integrase domains and the catalytic serine. Multiple sequence alignment was carried out with CLUSTAL Omega^46^ followed by manual editing in UGENE^47^. For clarity, protein sequence regions absent in ϕC31 were removed from the display. The positions of key residues are marked with coloured circles: red, the active serine; blue, residues forming the NTD dimer interface; grey, DNA-binding residues on αE; orange, residues forming the CC dimer interface; green, acidic hyperactive residues (dark green) and their interacting basic residues (light green); purple, RDF-interacting residues comprising acidic (light purple) and hydrophobic (dark purple) patches.

**Extended Data Fig. 4:**
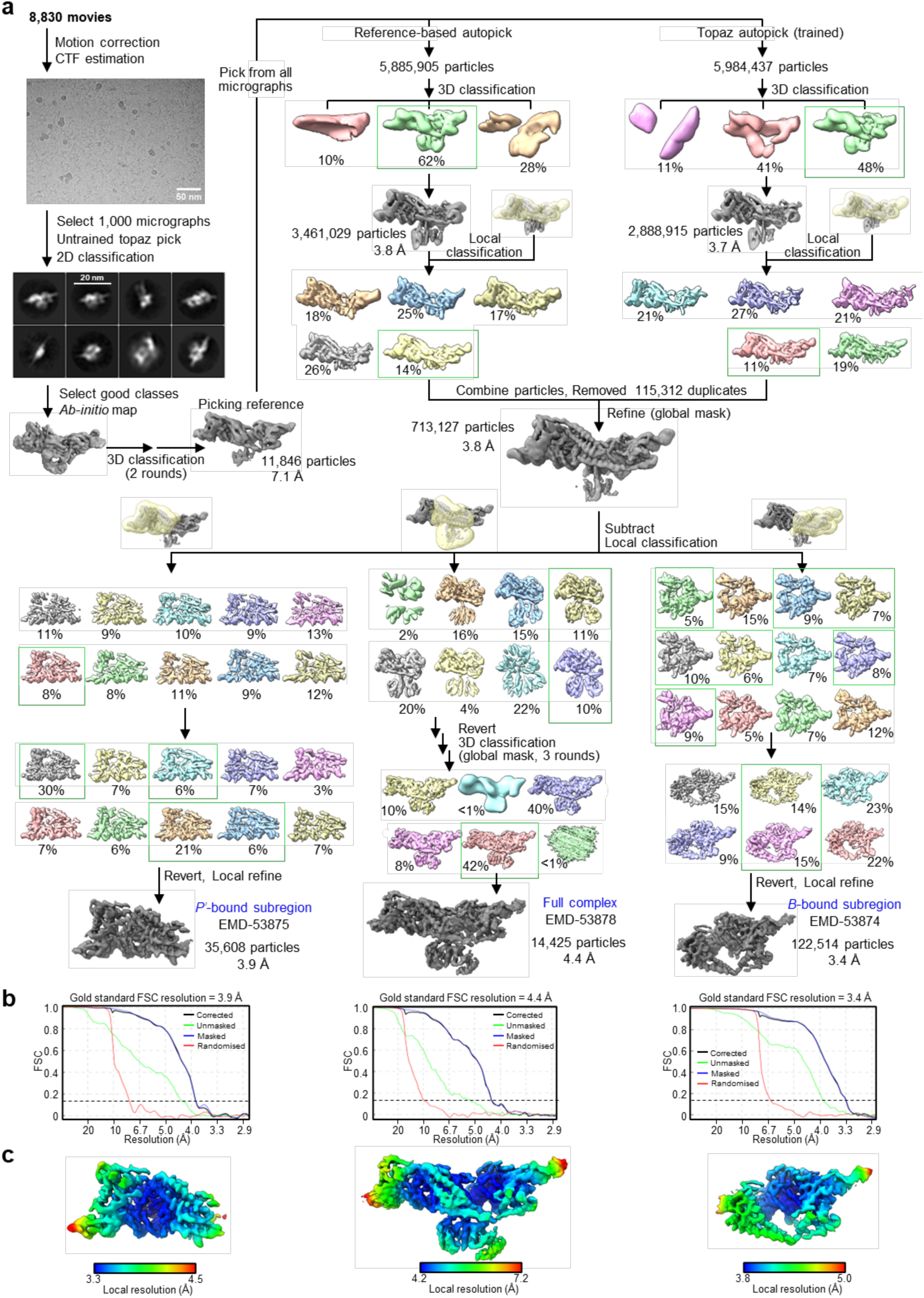
CryoEM analysis workflow and map quality metrics of the ϕC31 integrase–*attL* dimer complex. **a**, Cryo-EM image processing workflow. Particle classes selected for further processing are boxed in green, and final maps used for model building are labelled in blue. Masks used for local reconstruction, classification, or signal subtraction are shown as transparent yellow overlays. **b,c**, Gold-standard FCS curves used for global map resolution estimation (**b**) and local resolution estimate (**c**) of the final maps. Left to right: *B*-bound subregion; full complex; *P’*-bound subregion.

**Extended Data Fig. 5:**
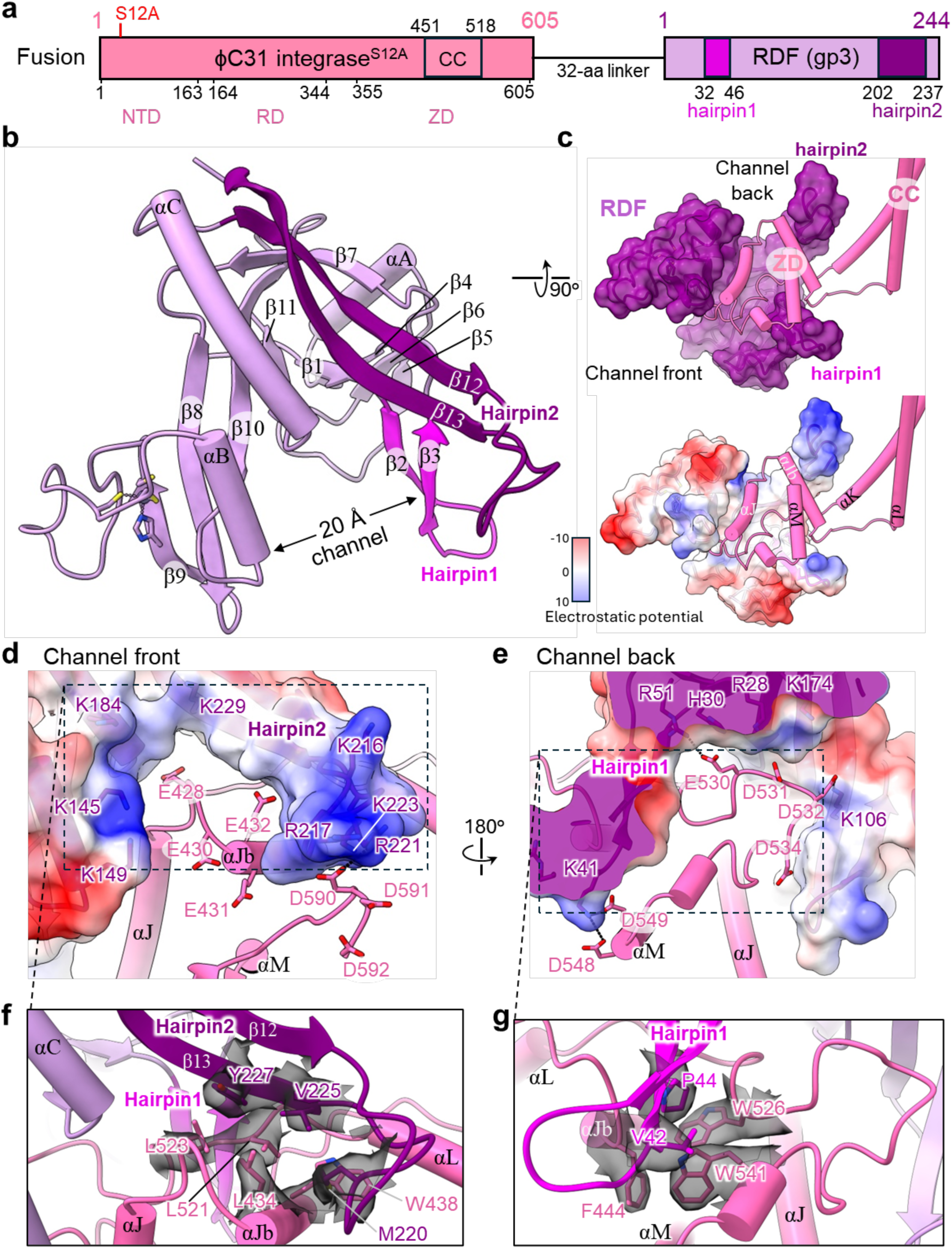
Details at the ZD–RDF interface. **a**, Details of the fusion integrase^S12A^-RDF construct used in cryo-EM experiments. The ϕC31 RDF, gp3 (UniProt ID Q9T216), is fused to the C-terminus of integrase^S12A^ with a 32-amino-acid glycine- and serine-rich linker. Protein features are shown and the boundaries are numbered according to their amino acid positions in each proteins. NTD, N-terminal catalytic domain; RD, recombinase domain, ZD, Zinc-ribbon domain; CC, coiled-coil motif; hairpin1 and hairpin2, two β-hairpin extentions that interact with the ZD tip and CC hinge. **b**, SEC-MALLS analysis of the complex formed by *attL* and the fusion protein. Elution profile is shown as solid line and the weight-averaged molar mass (M_W_) shown as circular markers. The expected mass of the dimer–*attL* and tetramer–*attL* complex (235 kDa and 470 kDa respectively) are marked with dash lines. **c**, Structure of RDF (monomer at the *P’* half-site in the fusion integrase^S12A^ -RDF–*attL* complex) viewed from the ‘front’ entrance of its ZD-binding channel. Hairpin1 and hairpin2 are in magenta and dark purple, respectively. Residues coordinating the Zn ion are shown as sticks. **d**, The ZD–RDF interface from an orthogonal view to that in **c** showing the length of the ZD-binding channel. Integrase is in pink. RDF is shown with a semi-transparent surface in purple (top) and coloured by electrostatic potential (bottom). **e**,**f**, Close-ups of the front (**e**) and back (**f**) entrances of the ZD-binding channel with integrase in pink and RDF in purple overlaid with surface electrostatics. Both sites feature interactions between acidic patches on ZD (including E428, D430, E431, and E432 at the αJ–αJb junction; D530, D531, D532, and D534 on the αL–αM loop; D548 and D549 on αM; and D590, D591, and D592 at the extreme C-terminus) and basic patches on RDF (K41 on hairpin1; K216, R217, R221, K223, and K229 on hairpin2; K145 and K149 on the large prong; and R28, H30, R51, K174, and K184 inside the channel). Residues forming these charged patches are shown as sticks. **g**,**h**, Close-ups of the hydrophobic interactions at hairpin2 (**g**) and hairpin1 (**h**). V42 and P44 on hairpin1 contact integrase’s αJb at F444 and the αL–αM loop at L521, L523, W526; M220, V225, and V227 on hairpin2 contacts integrase’s αJb at L434 and W438 and the αL–αM loop at L521 and L523. Hydrophobic side chains forming the interfaces are shown as sticks, with their cryo-EM densities overlaid in transparent grey.

**Extended Data Fig. 6:**
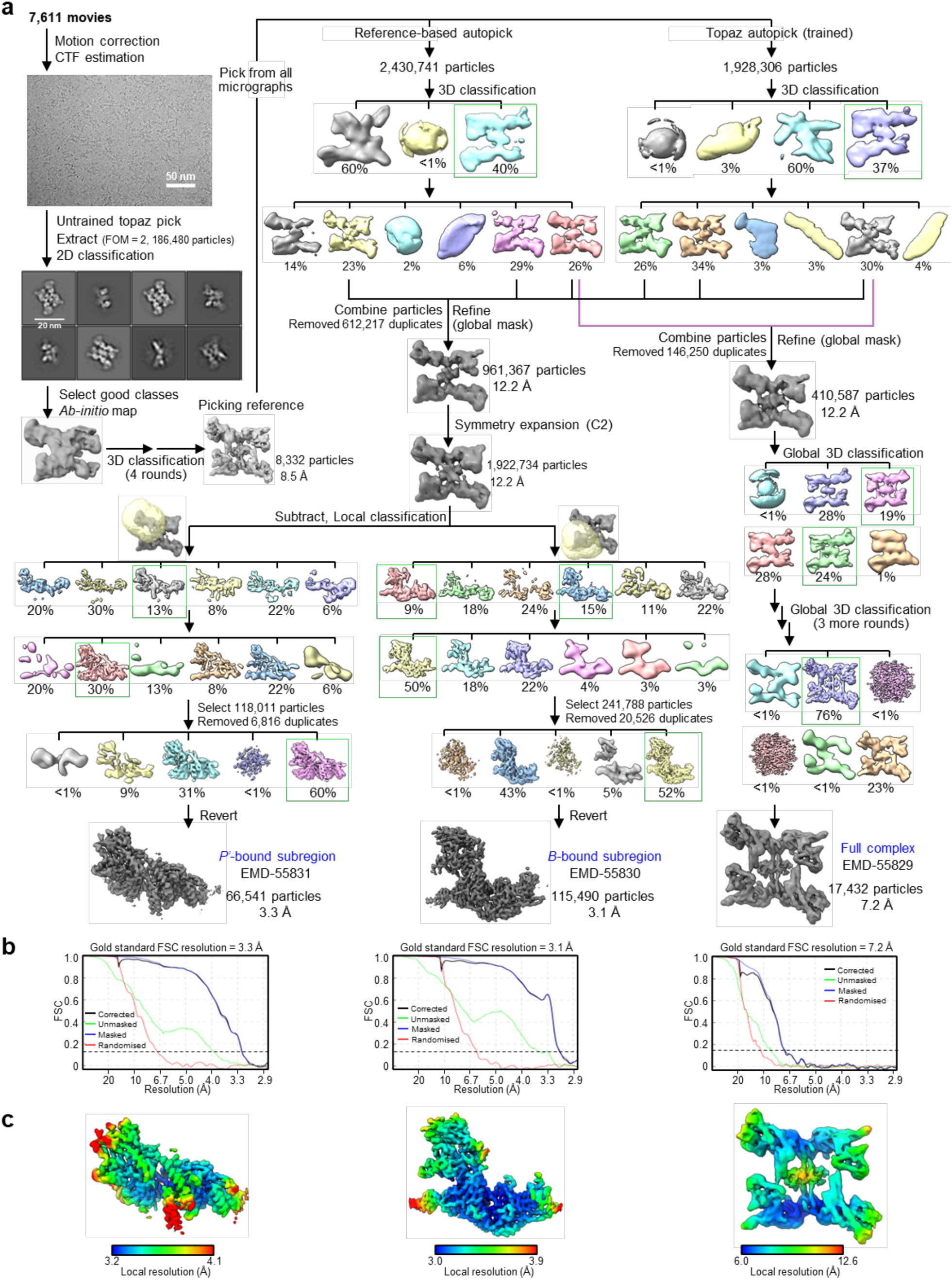
CryoEM analysis workflow and map quality metrics of the ϕC31 *attL*–*attL* synaptic complex. **a**, Cryo-EM image processing workflow. Particle classes selected for further processing are boxed in green, and final maps used for model building are labelled in blue. Masks used for local reconstruction, classification, or signal subtraction are shown as transparent yellow overlays. **b,c**, Gold-standard FCS curves used for global map resolution estimation (**b**) and local resolution estimate (**c**) of the final maps. Left to right: *P’*-bound subregion; *B*-bound subregion; full complex.

**Extended Data Fig. 7:**
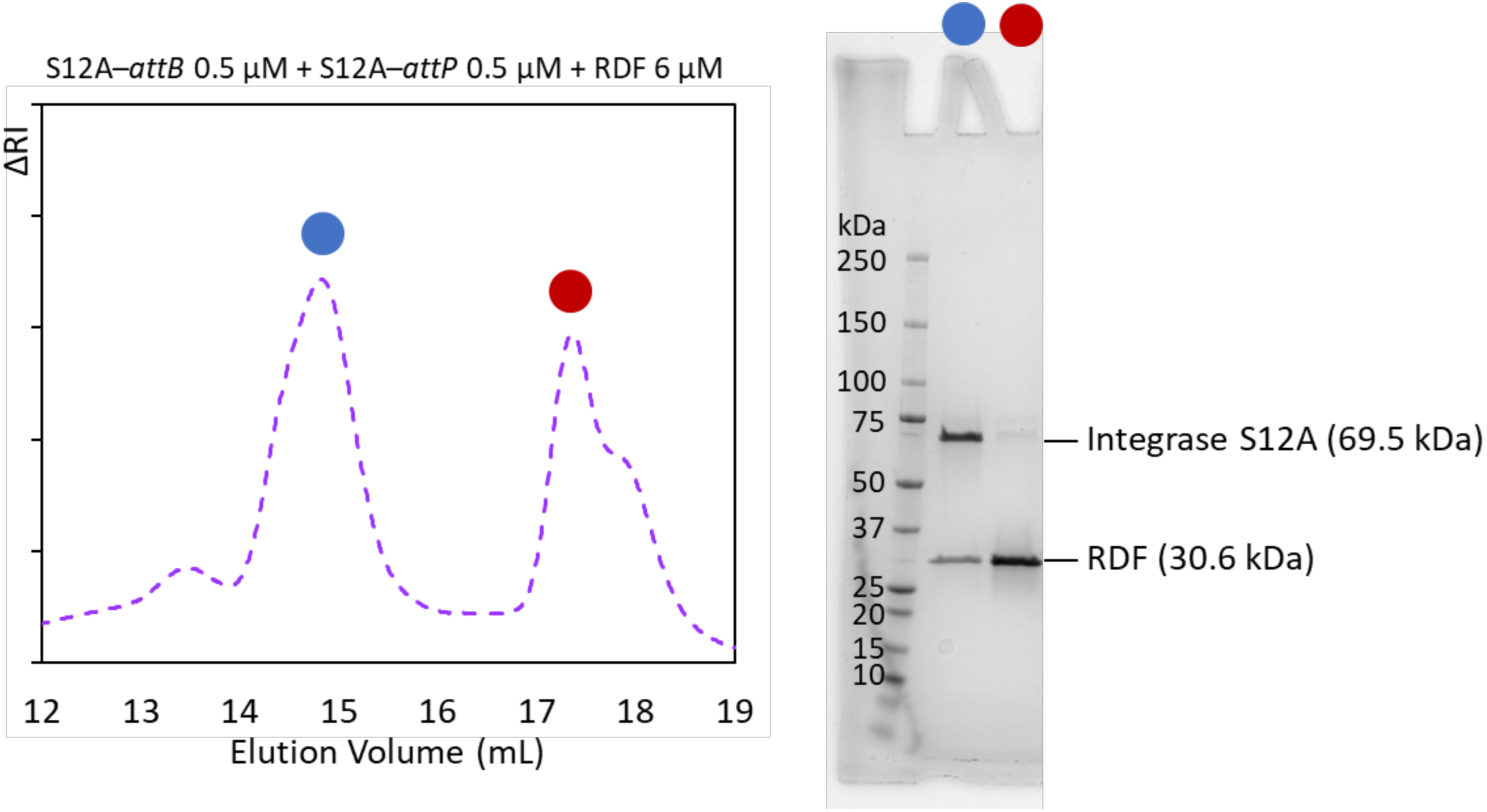
RDF induces attB–attP synaptic complex dissociation through direct binding. (Left) SEC analysis of a sample comprising the integraseS12A–*attB*–*attP* synaptic complex mixed with excess RDF, as shown in Fig. 4a. (Right) SDS-PAGE analysis of the two elution peaks at ∼15 ml and ∼17.5 ml, indicated in blue and red, respectively.

**Extended Data Fig. 8:**
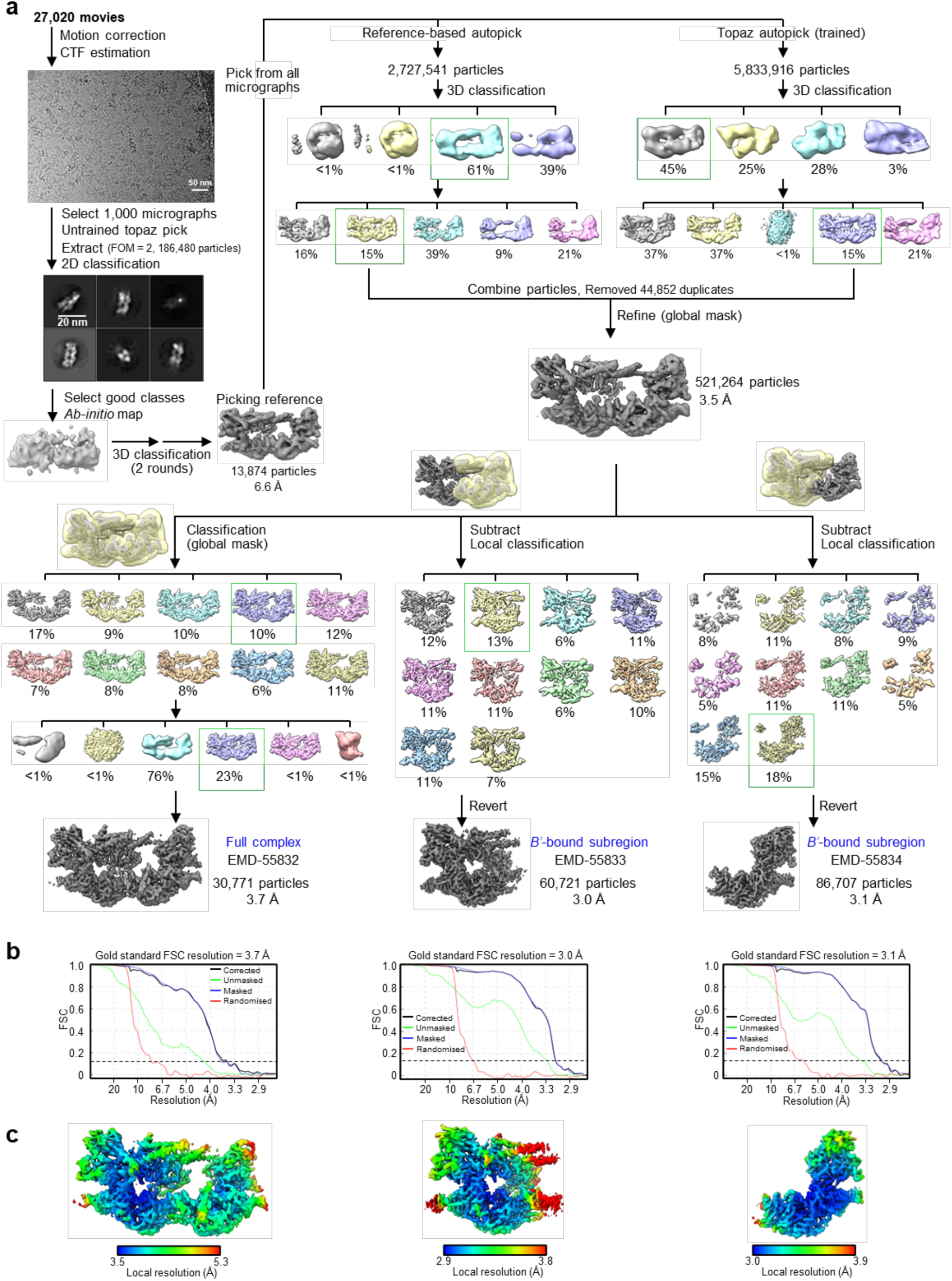
CryoEM analysis workflow and map quality metrics of the ϕC31 integrase^S12A^*– attB–*RDF dimer complex. **a**, Cryo-EM image processing workflow. Particle classes selected for further processing are boxed in green, and final maps used for model building are labelled in blue. Masks used for local reconstruction, classification, or signal subtraction are shown as transparent yellow overlays. **b,c**, Gold-standard FCS curves used for global map resolution estimation (**b**) and local resolution estimate (**c**) of the final maps. Left to right: full complex; *B*-bound subregion; *B’*-bound subregion.

**Extended Data Fig. 9:**
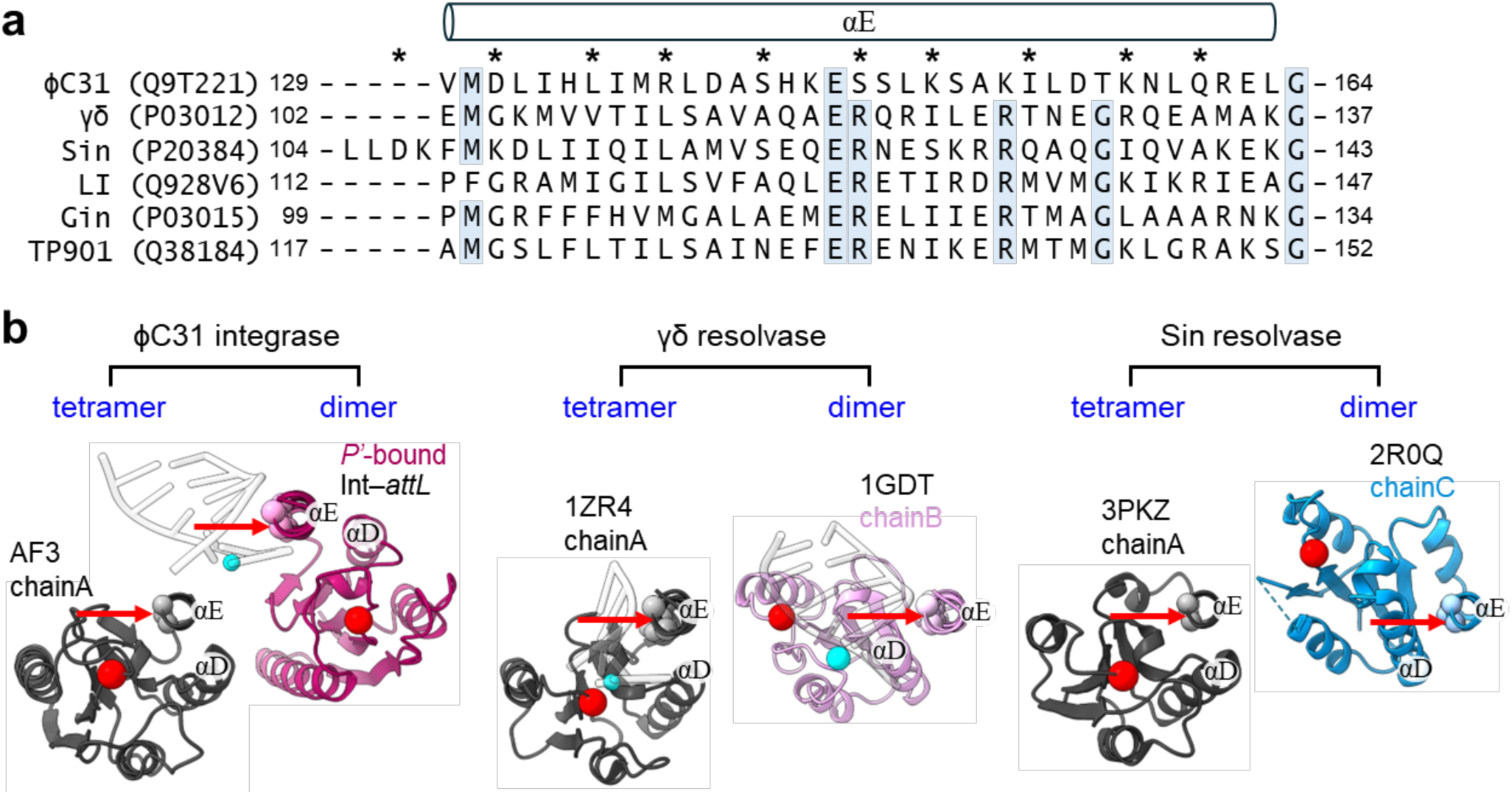
Trajectories of the NTD core region in dimer and tetramer structures of serine recombinases. **a**, Sequence alignment of the αE helix from selected serine recombinases with existing experimental structures. Protein names and Uniprot IDs (in parentheses) are shown on the left. Amino acid boundaries are indicated at the sequence ends, and those positions identical in five or more sequences are highlighted in blue. Residues on the DNA-binding face of αE are marked with an asterisk. The αE of Sin is one helical turn longer at the N-terminus. **b**, Structural comparison of the NTD monomer conformations within tetrameric and dimeric NTD structures of serine recombinases; these include ϕC31 integrase–*attL* dimer complex (magenta), AF3-predicted ϕC31 NTD tetramer (grey), γδ resolvase pre-cleavage dimer complex (PDB: 1GDT; pink), γδ resolvase post-cleavage tetramer synaptic complex (1ZR4; grey), Sin resolvase pre-cleavage dimer complex (2P0Q; blue), and DNA-free Sin resolvase NTD tetramer (3PKZ; grey). To show the variation in NTD core conformation relative to the αE helix, structures show αE helices in a similar orientation. The DNA-binding face of αE indicated by a red arrow; the Cα atoms of residues on this face (those asterisked in **a**) are shown as spheres (coloured according to the αE helices). Cα atoms of the active site serine residues (S12 for ϕC31, S10 for γδ, and S9 for Sin) are shown as red spheres. To show the relative positions of the scissile phosphates, base-pairs near the αE helix in the integrase–*attL*, 1ZR4, and 1GDT structures are shown in transparent white, with the scissile phosphorus atoms shown as cyan spheres. This analysis shows that the conformation of the ϕC31 NTD monomer in the AF3-predicted tetramer is similar to those in the tetramer structures of γδ and Sin; in this conformation, the active site serine residue is aligned to the DNA-binding face of the αE helix and is positioned adjacent to the scissile phosphate.

